# Site-specific metabolic labeling reveals proteome-wide phospho-dynamics

**DOI:** 10.1101/2024.07.23.604744

**Authors:** Mihai Alevra, Miguel Correa Marrero, Verena Kluever, Sunit Mandad, Nisha Hemandhar-Kumar, Kuan-Ting Pan, Julian van Gerwen, Selda Kabatas Glowacki, Hanna Wildhagen, Till Ischebeck, Yansheng Liu, Silvio O. Rizzoli, Henning Urlaub, Pedro Beltrao, Eugenio F. Fornasiero

**Author notes:** Equally contributing authors (Mihai Alevra, Miguel Correa Marrero, Verena Kluever, Sunit Mandad, Nisha Hemandhar-Kumar). Lead author, Twitter / X: @euforna (Eugenio F. Fornasiero).

## Abstract

Protein phosphorylation is a critical post-translational modification that orchestrates cellular signaling. Here, we introduce PulsPhos, a method combining metabolic labeling with phosphoproteomics, spectral analysis and modeling, to quantify site-specific phosphorylation lifetimes in living cells. Phosphosite lifetimes vary over multiple orders of magnitude and are influenced by factors such as amino acid composition and subcellular localization. PulsPhos was readily applied to pharmacological perturbations revealing fundamental mechanisms governing protein phosphorylation dynamics.

## Main

Advances in site-specific analysis of protein phosphorylation through mass spectrometry (MS)-based proteomics have shaped our understanding of key biological mechanisms, including temporal regulation of cellular processes such as cell cycle and cell signaling and enzyme activity, protein structure, localization, protein-protein interactions, and metabolism^1–5^. Work in this field has focused on uncovering changes in the relative abundances of protein phosphorylation in different conditions, or at different time points within specific processes, resulting in extensive catalogues of phosphorylation events^6–9^ and phosphorylation cascades^10^.

One of the main assumptions of the field is that phosphorylation sites have a limited lifetime. Despite the importance of this aspect, the lifetime values remain unknown. Due to the regulated action of kinases and phosphatases, phospho-groups are thought to undergo cycles of phosphorylation and dephosphorylation, at different rates. Depending on the nature of the biological process regulating (de)phosphorylation, the rates of exchange will change under different regimes. This makes the rate of phosphosite turnover a noteworthy aspect to study for understanding some of the basic principles shaping the regulation of the proteome. Their analysis has been attempted in *in vitro* preparations, as isolated nuclei^11^, or by using radioactive isotopes^12^, but there are currently no workflows that would allow the large-scale measurement of the phospho-group exchange rates in living cells.

Here, we have developed a safe, non-radioactive, experimental workflow, followed by innovative analytical tools compatible with conventional MS-based phosphoproteomics, that allow us to fill this gap, by estimating the average phosphorylation stability of phosphosites at the systems-scale level. Importantly, this approach should not be confused with those measuring the turnover of the phosphorylated proteins and peptides, which has been addressed in previous work^13,14^. Our simple, but effective approach uniquely uses a stable oxygen isotope (^18^O) to directly access the stability of phospho-groups and address phosphosite turnover. We have also established and validated all the experimental and data analysis tools and scripts required for the <u>p</u>roteome-wide <u>u</u>nveiling of labeling <u>s</u>tability in <u>phos</u>phosites (PulsPhos, **Fig. 1a**).

**Fig. 1:**
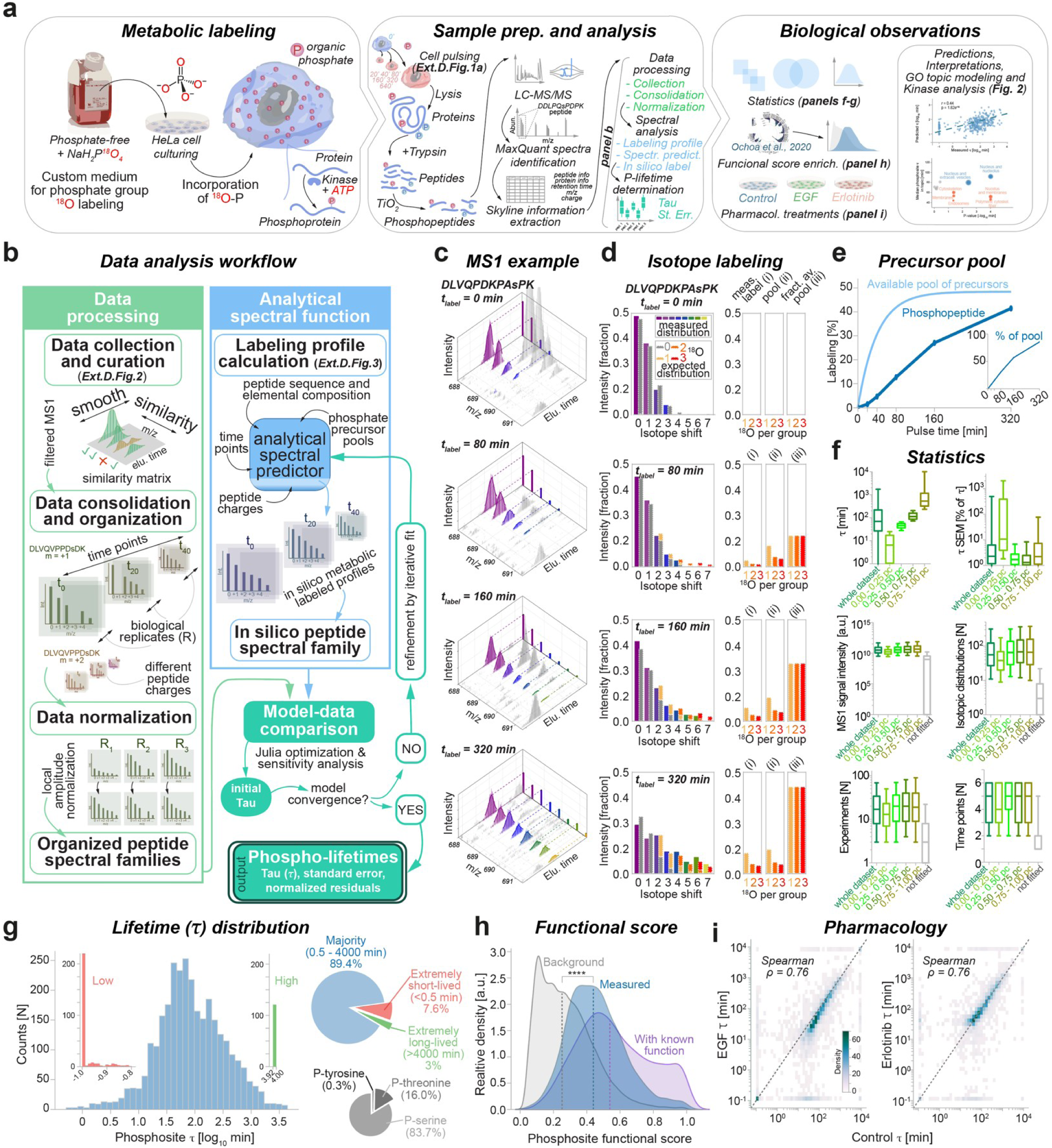
Determination of protein phosphorylation exchange rates with PulsPhos. . (**a**) The PulsPhos pipeline including metabolic labeling, sample preparation and analysis and biological data interpretation (see also Ext. Data Fig. 1). (**b**) Data analysis workflow (see also Ext. Data Figs 2-3). (**c**) MS1 labeling profile for one exemplary phospho-serine peptide (DLVQPDKPA<u>s</u>PK) over a 320 min. pulse. The m/z position is represented in rainbow scale. (**d**) MS1 isotope labeling profile measured from the same peptide (left panels, rainbow scale as in c), expected ^18^O distribution (left panels, gray/orange scale) and measured precursor label (measured distribution, rainbow scale), pool (expected distribution, orange scale), and fraction of the available pool (right panels, same orange scale). (**e**) Measured pool of precursor molecules (see also Ext. Data Fig. 1b-d) and exemplary labeling of a peptide. Note the sigmoid behavior of the peptide labeling indicative of a pool of precursors. Inset: peptide labeling expressed as percentage of available pool. (**f**) Statistics of median lifetimes expressed as Tau (*τ*), standard error of the mean (SEM), and other inputs subdivided among percentiles (pc) of *τ*, including excluded peptides during the consolidation step (not fitted). (**g**) Distribution of lifetimes, including classification (insets in color) and percentage of different phosphor-amino acids (inset in grayscale). (**h**) Functional score density of phosphosites with defined lifetimes vs. phosphosites with no function, and peptides with annotated function. (**i**) Comparison of lifetimes measured in a control situation with those measured following epidermal growth factor stimulation (EGF) or EGF receptor inhibition (erlotinib).

To achieve the metabolic labeling of protein phospho-groups in living HeLa cells, we developed a custom medium in which the naturally occurring ^16^O isotope in the monosodium phosphate (NaH_2_PO_4_) from the cell medium was replaced with stable ^18^O isotopes. The resulting labeled monosodium phosphate (NaH_2_P^18^O_4_) is non-radioactive, stable, and enables pulsed metabolic labeling of phosphate groups in living cells. This is then followed by phosphopeptide enrichment and liquid chromatography-mass spectrometry (LC-MS/MS) analysis (**Ext. Data Fig. 1a**). To obtain meaningful interpretations of the metabolic labeling with this pulsed approach, we determined the labeling status in the pool of precursor molecules (phosphate groups)^15^.

Labeling with ^18^O is non-radioactive, but in cells biotransformations progressively decrease the ^18^O content of the newly introduced phosphate groups, through the exchange with the ^16^O atoms of water. To account for these biotransformations, we conceived a strategy that enabled us to measure the ^18^O-^16^O exchange (**Ext. Data Fig. 1b-d**) and effectively allows us to obtain the availability of the pool of precursors necessary for the correct measurement of phosphosite lifetimes, here expressed as tau (τ; **Ext. Data Fig. 1e-h**).

Our data analysis workflow (**Fig. 1b**), comprises several steps, including data collection, curation, and consolidation (**Ext. Data Fig. 2**), as well as the *in silico* prediction of MS1 spectral families (**Ext. Data Fig. 3**), which is necessary for the determination of phosphosite lifetimes (see online Methods for details). Note that the metabolic labeling of phosphate groups, at increasing pulsing times, is evident in the MS1 spectra as a shift in the m/z values (**Fig. 1c**), while it is absent from non-phosphorylated peptides (**Ext. Data Fig. 4**), indicating the specificity of the ^18^O-phospate metabolic labeling. Moreover, the positive shift in the MS1 that we observe closely resembles the one that we can predict *in silico* based on the peptide amino-acid composition, the natural isotope abundance, and the availability of different ^18^O-phosphate precursors measured within cells (**Fig. 1d**). This is also consistent with the sigmoidal labeling behavior of individual phosphopeptides, confirming that the phosphosite exchange dynamics have a mono-exponential behavior with respect to the available pool of precursor molecules (**Fig. 1e**). Evaluation of the original input data in relation to the lifetime results shows that our data curation approach robustly avoids providing fitting results for poorly measured phosphopeptides (**Fig. 1f**).

**Fig. 2:**
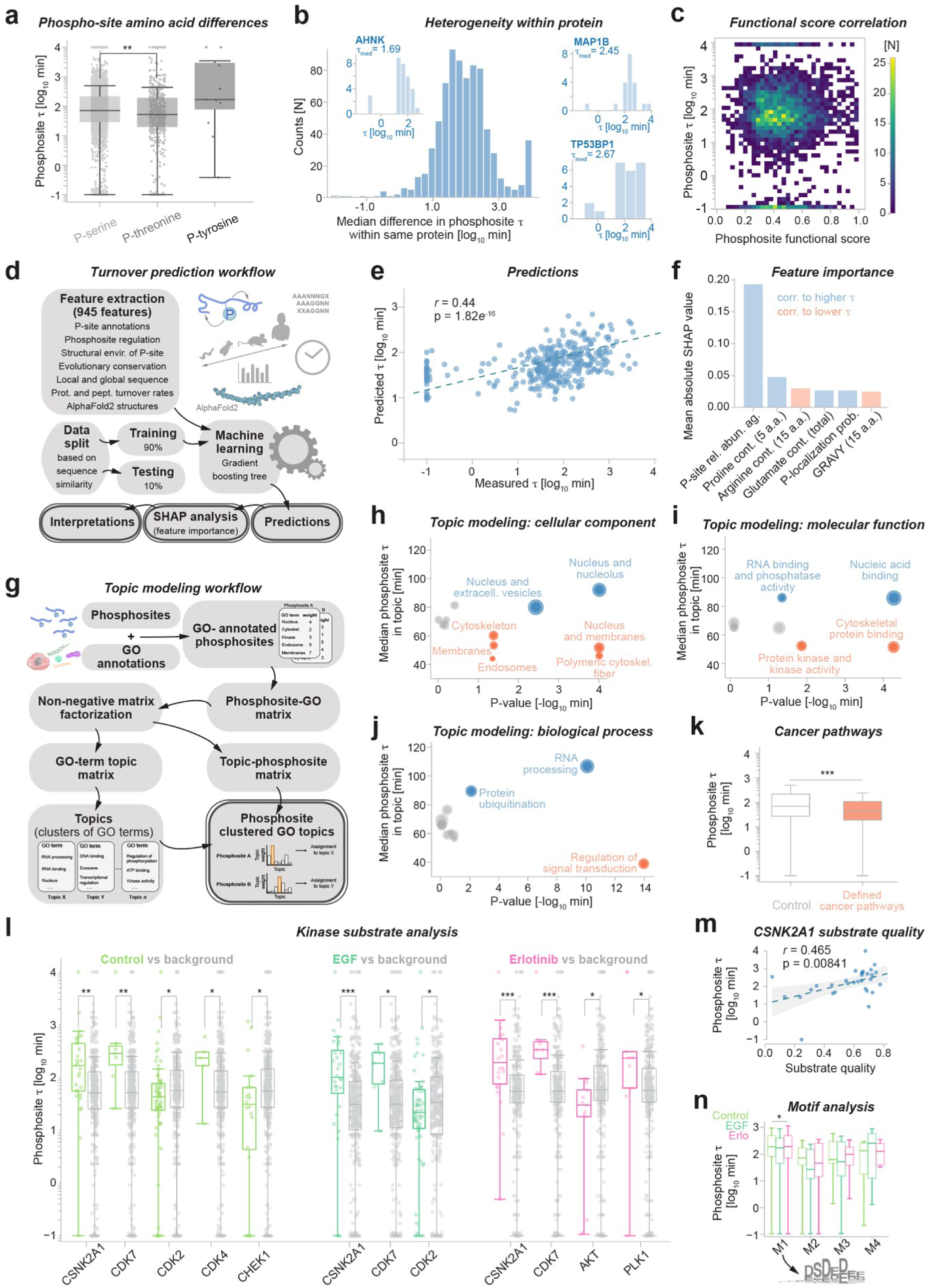
Measuring phosphorylation exchange rates reveals basic principles of phosphosite stability. (**a**) Boxplots of lifetime distributions subdivided by phospho-amino acid types. ** adjusted P-value <0.01 following Kruskal-Wallis test (p=0.0015) (**b**) Lifetime heterogeneity calculated “at the whole protein level” for proteins with more than one phospho-peptide, including examples for proteins with different median lifetimes (insets). (**c**) Lifetime vs. functional score correlation showing equal importance of phosphopeptides regardless of differences in lifetimes. (**d**) Workflow used for the prediction of lifetimes (see online methods for details). (**e**) Machine learning predictions on the test set. (**f**) Feature importance analysis for the predictions shown in e and correlation to either higher lifetime (slower turnover) or lower lifetime (higher turnover). (**g**) Topic modeling workflow (see online methods for details) and results for different gene ontologies (GOs; **h-j**). Dot size is proportional to the number of phosphosites in the topic, red indicates *τ* within the topic higher than the median, blue lower *τ*, and gray no difference. Refer to Ext. Data Fig. 6a-e for additional analyses (**k**) Lifetime analysis for cancer pathways. (**l**) Kinase substrate analysis for different treatments revealing an association between kinases and lifetimes. Kolmogorov-Smirnov test followed by Benjamini-Hochberg correction (**m**) Substrate quality for the casein kinase 2 (CSNK2A1) correlates positively with lifetimes (linear regression with 95% confidence intervals is shown). (**n**) Phospho-motif analysis identifies a putative CSNK2A1 motif positively associated with larger lifetimes (aspartic and glutamic acid rich, reported in the gray inset below the graph).

The phosphosite lifetime distribution measured at the whole proteome level reveals that the median time of phospho-group exchange on phosphopeptides is ∼70 minutes in HeLa cells. Only ∼7.5% of these events lasts <30 seconds, while ∼3% are extremely stable, lasting >33h (**Fig. 1g**). The sites for which we measured the phosphosite lifetime in our dataset have significantly higher functional scores ^7^ than the background, constituted by all identified phosphosites (**Fig 1h**), reflecting their role in protein function and suggesting their importance for organismal fitness.

We also performed measurements under different pharmacological conditions that alter EGF signaling, which indicate that phosphosite lifetime measurements are reproducible across cellular modulations (**Fig. 1i**), similarly to what it observed for other proteome features such as protein abundance (**Ext. Data Fig. 5**).

A thorough analysis of our results revealed several intriguing biological observations (**Fig. 2** and **Ext. Data Fig. 6, Ext. Data Table 1**), underscoring the significance and applicability of PulsPhos to future research challenges. For instance, we observe that serine phosphosites are more stable than threonine phosphosites (**Fig. 2a**). Additionally, we found extensive variability of lifetimes of sites found within the same protein, spanning orders of magnitude (**Fig. 2b**), pointing to dissimilar regulation of different phosphosites.

The lack of correlation between lifetimes and functional scores (**Fig. 2c**) indicates that the importance of phosphorylation events for cellular fitness is not skewed towards very short or very long lifetimes. We carried out a three-pronged analysis to study the determinants of phospho-lifetimes. First, we trained a machine learning model to predict he average lifetime of phosphorylation events (τ), based on a broad set of features (**Fig. 2d**), whose predictions correlate reasonably and significantly with the measurements (**Fig. 2e**). Amongst the most important features, we found that an aggregated feature for protein abundance and phosphosite occupancy, local proline, and total glutamate content correlate to a longer τ, while local arginine content and local hydrophobicity correlate to a shorter τ (**Fig. 2f**). Moreover, gene ontology (GO) topic modelling, pathway analysis (**Fig. 2g-k**) and gene set enrichment analysis (**Ext. Data Fig. 6**) indicate that processes occurring in the nucleus are linked to slower phosphosite exchange rates, while signal transduction and protein kinase activity are linked to faster exchange rates. Finally, kinase substrate and phosphorylation motif analysis revealed general tendencies of specific kinases to be associated to more stable phosphorylation events, such as the casein kinase 2 alpha 1 (CSNK2A1; **Fig. 2l-n** and **Ext. Data Fig. 6g**), and that the erlotinib block of EGF receptor signaling is able to slow down several biological processes in a phosphorylation-dependent manner (**Ext. Data Fig. 6i**).

Overall, with PulsPhos we provide the community with a novel method to address a dynamic aspect of protein phosphorylation regulation that was previously inaccessible, allowing the direct study of the dynamics of phosphorylation processes in cells. We found that the lifetimes of phosphorylation sites vary widely, with phospho-serine sites generally more stable than phospho-threonine sites. These lifetimes are influenced by factors such as amino acid content, local hydrophobicity and subcellular compartmentalization, and are tailored for specific functions (e.g., faster for signal transduction but slower for RNA processing) and influenced by the activity of specific kinases (as in the case of CSNK2A1). By introducing site-specific analysis of phosphorylation flux and enabling its combination with other approaches to study the dynamic stability of the proteome ^16^, PulsPhos promises to deepen our understanding of the fundamental principles of proteome regulation.

## Online Methods

### Chemical synthesis of NaH_2_P^18^O_4_

For the study, we synthesized ^18^O-labeled phosphoric acid by using phosphorus pentachloride and 97% water-^18^O ^17^. Phosphoric acid-^18^O_4_ was neutralized by a solution of sodium hydroxide and precipitated with ethanol to give crystalline sodium phosphate-^18^O_4_.

#### Reagents

Phosphorus pentachloride and 97% water-^18^O were purchased from Merck (former Sigma Aldrich, Darmstadt, Germany). Toluene and ethanol were obtained from commercially available sources with grade “*puriss. p.a.”* and were used as supplied. Sodium hydroxide was delivered from Carl Roth (Karlsruhe, Germany). Demineralized water for the preparation of sodium hydroxide solution was purified prior use with MilliQ Advantage A10 from Merck Millipore (Darmstadt, Germany). Deuterated water for NMR was purchased from Deutero (Kastellaun, Germany).

#### Reactions

Phosphorus pentachloride was weighed in a glove box and stored under vacuum until usage. The air- and water-sensitive reaction was conducted under inert atmosphere. Therefore, glass equipment was flame-dried under vacuum followed by applying a purge-and-refill technique under argon atmosphere.

#### Instruments

The ^31^P-NMR spectrum was measured on a Bruker Avance III 400 (Rheinstetten, Germany). Chemical shifts are quoted in ppm (TMS = 0 ppm). Multiplicities are abbreviated as follows: s = singlet and br s = broad singlet. Freeze-drying of compound from aqueous solution was performed using a Christ-Alpha-2-4 lyophilizer attached to a high vacuum pump and a Christ RCV-2-18 ultracentrifuge (Osterode am Harz, Germany).

#### Synthesis

Phosphorus pentachloride (2.35 g, 11.3 mmol, 1.0 eq.) was cooled down with liquid nitrogen followed by dropwise addition of heavy water (0.90 mL, 1.00 g, 50.0 mmol, 4.4 eq.). The reaction mixture was slowly warmed to room temperature (RT), stirred at 80 °C for 90 min and then bought to RT. The resulting phosphoric acid-^18^O_4_ was first co-evaporated with toluene (3 *x* 5 mL), then adjusted with aq. NaOH (1 M) to a pH of 7.4−¿7.8 and finally freeze-dried. The white solid was suspended in ethanol (25 mL), filtered, washed with ethanol (3 *x* 5 mL) and dried *in vacuo*. The product, sodium phosphate-^18^O_4_, was confirmed with ^31^P-NMR (162 MHz, D_2_O, RT): δ = 2.26−¿2.32 (br s, HPO_4_^2-^, H_2_PO_4_^-^) ppm.

### Cell culturing

HeLa cells (Obtained from Leibniz Institute DSMZ-German Collection of Microorganisms and Cell Cultures DSMZ no.: ACC 57) were grown to passage five in Dulbecco’s Modified Eagle Medium (DMEM; Sigma) supplemented with 10% FCS (Gibco), 2mM L-Glutamine (ThermoFisher), 100 U/mL penicillin (Sigma) and 100 mg/mL streptomycin (Sigma) at 37°C, 5% CO_2_. Cells were split for optimal growth four and one day prior to the experiment and were 80% confluent at the endpoint of the time course. For GC-MS measurements, cells were plated in 60mm dishes (VWR), for LC-MS, cells were plated in 145mm dishes (Greiner Cellstar).

### Cell media and buffers for both labeling and 2-Deoxy-D-Glucose experiments

The custom ortho-phosphate medium, pH 7.4, was prepared fresh on the day of the experiment according as reported below. Cells tolerate the medium as it is very close to a standard MEM medium, in which the osmolarity id adjusted with NaCl based upon addition of NaH_2_P^18^O_4_ (see below). After addition of NaH_2_P^18^O_4_, the osmolality was adjusted to 365 mOsm with ddH_2_O and 5M NaCl using a Gonotec Osmomat 030. The medium was then filtered for sterile use. For the iso-osmolar wash buffer, 9.6g NaCl were dissolved in 900mL ddH_2_O to result in a 365 mOsm solution. For the 2-Deoxy-D-Glucose (2DG) experiments, 1025mg of powdered 2-Deoxy-D-Glucose (D8375 Sigma) were dissolved in 25mL DMEM, high glucose, no phosphates (11971025 Gibco/Thermo). Before addition of the ^18^O-ortho-phosphate medium (2.5mL for 60mm dishes and 10mL for 145mm dishes), cells were washed three times with iso-osmolar wash buffer. 5 minutes before collection of the cells, 2-Deoxy-D-Glucose was spiked into the medium, see **Extended Data Fig. 1b-c** for a detailed explanation of its use for the pool labeling analysis. For collection, cells were washed quickly three times in iso-osmolar wash buffer before addition of either extraction buffer for GC-MS or lysis buffer for LC-MS and subsequent scraping. Samples for GC were kept at -20°C and MS samples were snap frozen in liquid nitrogen and stored at -80°C until analysis.

### ^18^O-ortho-phosphate medium composition for 500 ml

| Amount | Ingredient | Cat. No. |
| --- | --- | --- |
| To final 0.5L volume | ddH<sub>2</sub>O | n.a. |
| 100mg | CaCl<sub>2</sub> | Sigma C1016 |
| 0.05mg | Ferric Nitrate Fe(NO<sub>3</sub>)<sub>3</sub>°9H<sub>2</sub>O | Sigma 254223 |
| 49mg | MgSO<sub>4</sub> | Sigma M2643 |
| 200mg | KCl | Sigma P5405 |
| 1850mg | NaHCO<sub>3</sub> | Sigma S5761 |
| 2250mg | D-Glucose | Sigma G7021 |
| 15mg | Glycine | Sigma G5417 |
| 21mg | Serine | Sigma S4311 |
| 20mL | MEM Amino Acids (50x) solution | Sigma M5550 |
| 20mL | MEM Vitamin Solution (x100) | Thermo 11120052 |
| 10mL | L-Glutamine | Thermo 25030081 |
| 50mL | FBS dialyzed | Gibco A3382001 |
| 7.231g | NaH<sub>2</sub>P<sup>18</sup>O<sub>4</sub> | Custom synthesis |
| To final 365 mOsm | NaCl 5M | Sigma 59222C |

### Pharmacological treatments

Samples were treated either with hEGF (E9644 Sigma; 20ng/mL final) or with Erlotinib HCl (S1023 Absource Diagnostics; 10µM final) 12h before the start of the experiment and directly again directly after the change to the ^18^O-ortho-phosphate medium until the end of the experiment. The hEGF powder reconstituted in 10mM acetic acid (Roth, 99%, 60.05g/mol #7332) at 1mg/ml stock which was diluted in the custom medium.

### Determination of the available pools of labeled phosphates by GC-MS

Cells were washed and resuspended in 300 µl organic solvent (methanol/chloroform/water 32.25:12.5:6.25 [v/v/v]) thereby extracting its metabolites. Phase separation was induced with 150 µl of 0.05 mg/ml *allo*-inositol in water. The mixture was vortexed and centrifuged for 5 minutes at 2,000 g and 4°C. 150 µl of the upper polar phase was dried under nitrogen stream and derivatized with 10 µl methoxyamine hydrochloride and 20 µl N-methyl-N-(trimethylsilyl)trifluoroacetamide (MSTFA) as previously described^18^ to transform the metabolites into their methoxyimino (MEOX)- and trimethylsilyl (TMS)-derivatives.

Samples were analyzed on an Agilent 5973 Network mass selective detector connected to an Agilent 6890 gas chromatograph equipped with a capillary HP5-MS column (30 m x 0.25 mm; 0.25 µm coating thickness; J&W Scientific, Agilent). Helium was used as carrier gas (1 ml/min). The inlet temperature was set to 230 °C and the temperature gradient applied was 50 °C for 2 min, 50-330 °C at 15 K/min, and 330 °C for 2 min. Electron energy of 70 eV, an ion source temperature of 230 °C, and a transfer line temperature of 330 °C was used. Spectra were recorded in the range of 71-600 Da/e. 2 µl of sample were injected.

For quantification of the isotopic distribution in the phosphate group of 2-Deoxy-D-Glucose-phosphate, the fragment with the monoisotopic mass-to-charge ratios 299 Da/e was analyzed as it represents only the phosphate group of the molecule and its TMS-groups from the derivatization. Non-labeled samples were used to determine the natural isotope background of the fragment. This background was taken into consideration for the calculation of the ^18^O labeling.

### Mass spectrometry sample preparation

After washing HeLa cells quickly with the iso-osmolar wash buffer three times each petri dish (corresponding to ∼2 mg of total protein content) was quickly lysed in 1 ml of lysis buffer (8 M Urea, 200 mM HEPES, 20 mM Tris (2-carboxyethyl)phosphine-hydrochloride (TCEP), 80 mM 2-Chloroacetamide (CAA), 1 × Halt Protease inhibitor (ThermoFisher Scientific) and phosphatase inhibitor (ThermoFisher Scientific) cocktails) and quickly frozen in liquid nitrogen to block possible changes in phosphorylation. For downstream protein purification, lysates were incubated at 95°C. After heating the samples, they were cooled on ice and they were sonicated for 5 min using 30 seconds ON/OFF cycles with a Bioruptor ultrasonication device (Diagenode, Seraing, Belgium) that was used at maximum duty output. After sonication, proteins were precipitated by adding 80% acetone and incubated overnight at -20°C. The next day samples were centrifuged at maximum speed (∼20’000g) in a tabletop centrifuge (Eppendorf) for 60 min. The supernatant was discarded, and the pellet was washed with absolute acetone and maximum speed (∼20’000g) in a tabletop centrifuge (Eppendorf) for 10 min. After discarding the supernatant, the protein pellets were air dried for 20 min and resuspended in 200 µl of 200 mM HEPES, 20 mM TCEP, 80 mM CAA supplemented with protease inhibitor and phosphatase inhibitor cocktails. To resuspend the pellets, samples were sonicated once more for 5 min using 30 seconds ON/OFF cycles on the same Bioruptor ultrasonication device. To ensure perfect processing, samples were then incubated for 30 min in a thermomixer at 37°C 750 rpm. After alkylation, 20µg of mass spectrometry grade Trypsin gold (Promega) was added to each tube and proteins were digested overnight at 37 °C.

The next day samples were desalted using prepacked C18 spin columns (Harvard Apparatus, Holliston, USA) and further processed to enrich phosphopeptides with TiO_2_ beads (GL Sciences, Tokyo, Japan) at 1:10 protein-to-beads ratio for 20 min at 40°C. The unbound peptide fraction (nonphosphorylated peptides) was discarded, and TiO_2_ beads were thoroughly washed four times in washing buffer (60% (v/v) acetonitrile (CAN), 1% (v/v) trifluoroacetic acid (TFA) in water). The phosphopeptides that were bound to beads were eluted using 3.75% (v/v) NH_4_OH 40% (v/v) ACN in water, snap-frozen in liquid nitrogen, and dried in a centrifugal Savant SpeedVac vacuum concentrator (ThermoFisher Scientifi). Dried phosphopeptides were resuspended in 200 mM HEPES and desalted using the prepacked C18 spin columns. Finally, peptides were concentrated in a SpeedVac and stored at -20°C.

### Mass spectrometry sample analysis

The dried phosphopeptides were resuspended in 20 μl of sample loading buffer (5% (vol/vol) LiChrosolv-grade ACN and 0.1% (vol/vol) formic acid in LiChrosolv-grade water). Vials were sonicated in a water bath for 3 min (at maximum cycle time). Vials were centrifuged in a table-top centrifuge at 12’000g for 5 min at room temperature to avoid the debris. The supernatant was transferred to fresh LC-MS glass vials. Samples were automatically injected with an online UltiMate 3000 RSLCnano HPLC system coupled to Q Exactive HF for reverse phase LC-MS/MS.

Loaded peptides were injected in a reverse-phase C18 Acclaim PepMap100 5-μm trapping-column for 3 min. After 3 min, switch the valve online on an analytical C18 column (30-cm length, 75-μm i.d.; produced in-house using a ReproSil-Pur C18 AQ 1.9-μm reverse-phase resin). Fractionated peptides were eluted with buffer B (95% (vol/vol) LiChrosolv-grade ACN and 0.1% (vol/vol) formic acid in LiChrosolv-grade water), creating a gradient at a flow rate of 300 nl/min over a 118-min gradient time. The pre-column and column temperatures were kept at 50 °C during the whole chromatography. MS data were acquired by scanning the precursors in a mass range from 350 to 1,600 Da at a resolution of 60,000 at m/z 200. The 30 most intense precursor ions were chosen from MS1 for MS2 fragmentation. For MS2, higher-energy collisional dissociation (HCD) fragmentation was performed with the AGC target fill value of 1 × 10^6^ ions. The precursors were isolated with a window of 1.6 Da.

### Initial mass spectrometry data analysis

To identify the phosphopeptides, the .RAW files were analyzed in MaxQuant (version in 1.5.2.8) using default settings with phospho-STY included as variable modifications. No O^18^ was initially included as a parameter, since we developed an independent spectral analysis which accounts for this. The canonical amino acid sequences of *Homo sapiens* proteins were retrieved from the Uniprot database containing 71913 referenced, 20192 reviewed and 51721 unreviewed entries. Potential contaminants, reversed sequences, and phosphorylation sites identified with localization probability <0.75 were not further considered for analysis. Official gene names and Uniprot accession code of the leading protein were used in all subsequent analyses. The MaxQuant output phosphopeptide files were used as an input for Skyline version 3.6.0.11714 to consolidate the information necessary to perform the subsequent spectral analysis carried on with MATLAB and Julia (see below). In detail, the information gathered with Skyline used in downstream analyses corresponded to: 1) the peptide and protein information; 2) the retention time window and 3) the m/z from each single experiment. This allowed to match all the phosphopeptide identifications across experiments and replicates. To be able to analyze the files in MATLAB, we also converted the .RAW files with Thermo Proteome Discoverer 2.0 (Thermo Fisher Scientific) into the .MZ5 file format.

### Custom spectral analysis for metabolic labeling

We imported the .MZ5 files with a custom-made script in Julia (Version 1.9.2) to extract all raw spectra, transforming mass differences to absolute masses and saving individual (MS1) peaks that were separated by zero intensity in the spectra. The intensity-weighted center of mass for each retention time was determined for each peak (red vertical line in Extended Data Fig. 2c), and the similarity of each curve to a gaussian with identical standard deviation, amplitude and center was used as quality measure to reject non-gaussian, typically multi-component peaks. A theoretical isotopic mass distribution (including relative amplitudes *I*_n_ and mass centers *M*_n_ for the first eight isotopic positions from n=0 to n=7) for each detected peptide was calculated from its amino acid sequence using a custom-written Julia adaptation of the fast Poisson model as previously described^19^. In cases where multiple possible phosphosites were reported with individual detection probabilities for a peptide, only the most probable (detection probability > 0.75 of maximum probability) were selected in each experiment. Individual peak data (peak center masses and intensity) was then collected from the whole dataset for each such detected phosphosite, filtering at mass windows of 4 ppm around the first 8 theoretical isotopic mass centers *M*_n_ for each detected charge, and retention time windows of 0.75 s around the center retention time reported for each experiment. For each charge, experiment and isotopic peak number, an intensity histogram was calculated along the retention time and cross-correlated with that of the first isotopic number to remove non-similar retention time curves that were a typical result of contamination with overlapping spectra of other peptides. For the selected retention time curves, all peaks inside of one standard deviation from the center retention time were gaussian-smoothed and the maximum of the resulting curve saved as “spectral fingerprint” for later fitting.

For each treatment (control, EGF and erlotinib), the phosphorylation lifetime τ_p_ for each phosphosite was estimated by least-squares fitting of a metabolic labeling model to the isotopic “fingerprints” saved above, weighting the dataset of each experiment and charge by their respective sum of amplitudes and normalizing amplitudes along the available isotopic peak numbers to calculate the total residual.

The metabolic labeling model describes the temporal evolution of the isotopic mass distribution during labeling time *t*, assuming a mono-exponential change of available labeled phosphates (*y*_0_ … *y*_3_ for non-labeled to triple-labeled ^18^O phosphates) with a pool lifetime of τ, and a resulting double-exponential change of labeled phosphorylations (*p*_0_ … *p*_3_) following the available pool phosphates with an individual phosphorylation turnover time constant τ_p._

With a general normalized pool labeling intensity of*l* (*t*) =1−exp (−*t* / *τ*), non-labeled phosphates are then given as:

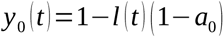

and labeled phosphates (i=1…3) as

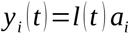

Individual equilibrium contributions *a*_i_ of the available phosphates were estimated using the measurements obtained from the 2-Deoxy-D-Glucose experiments (Extended Data Fig. 1b-d), as well as an initial guess of the pool time constant τ, using least-squares fitting. The four species of 0 to 3-labeled phosphorylations then follow the pool at a phosphosite-specific time constant τ_p_, with the overall isotopic labeling being:

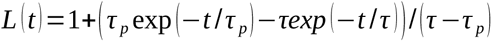

and individual (non-labeled and labeled) phosphorylations described by

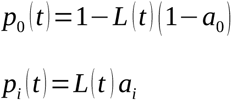

The combined predicted mass spectrum amplitude *A*_n_ at isotopic positions n and labeling time *t* is then calculated from the non-labeled reference spectrum *I*_n_ as weighted sum of the possible phosphorylations, each shifted by two neutron masses per additional ^18^O label:

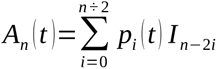

Residuals between the predicted and observed amplitudes for all phosphosites and phosphopeptides were minimized, giving individual phosphorylation turnover time constants τ_p_ and a confidence interval Δ τ_p_ from the fit process. For some peptides however, a lower bound (0.1 s) for τ_p_ was reached, signifying a labeling speed that was faster than the assumed labeling time constant τ for the pool of phosphates. To account for a possible availability of labeled phosphates that was faster than the measured proxy, we did a parameter sweep for τ, repeating the whole fit process and globally minimizing the sum of two cost functions (normalized to the initial fit): 1) the number of phosphopeptides that reached the lower border, 2) the sum of fit precision (Δτ_p_/τ_p_) for all phosphopeptides. The resulting pool time constant τ was then used for the final individual turnover time constants, as a compromise between underestimating the labeling speed of the pool and overall model precision.

### Dataset curation

The analyses described below were performed using a high-confidence subset of the whole data. First, phosphopeptides containing multiple phosphorylations were not considered in this first work, to avoid determination issues, since downstream analyses focus on properties of individual phosphosites. Next, fit results of τ_p_ with a coefficient of determination (R^2^):

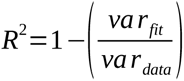

Values below 0 were discarded because of their poor fit (as negative R^2^ in a non-linear model are expected when the fit explains the data worse than the average value would, i.e. for data that shows a negative mass shift over labeling time, which is impossible within the constraints of our physical model). To further increase the reliability of the dataset, we used only phosphosites found in the high-confidence reference human phosphoproteome curated by Ochoa *et al.*^7^. In the case a phosphopeptide can be mapped to multiple proteins in UniProt, but it can unambiguously be assigned to one of the phosphosites in the reference phosphoproteome, we consider it to be this phosphosite. These filtering steps provided 4058 phosphosites in our final dataset (see Extended Data Table 1).

### Feature extraction

We gathered 945 features to describe phosphosites and subsequently train machine learning models. Amongst these, we included mass spectrometry evidence from our experiments (number of independent detections and number of peaks). We also used the phosphosite features computed by Ochoa *et al.* ^7^ for their curated dataset of high confidence phosphosites. These features comprise four broad categories: MS evidence for the phosphosite (*e.g.*, the peptide posterior empirical probability and spectral counts); phosphosite regulation (*e.g.*, matching kinase motifs and the kinase with highest significant co-regulation); the structural environment of the phosphosite (*e.g.*, sequence-based predictions of surface accessibility and disorder) and evolutionary conservation (*e.g.*, SIFT scores and phosphosite evolutionary age). We refer the readers to the original manuscript^7^ for more precise information about these features.

In addition to these precomputed features, we extracted additional features from protein sequences to describe both the whole protein and the region around the phosphosite (in the latter, using windows of 5, 10, 15 and 20 residues around the phosphosite). These features, both global and local, include the isoelectric point, net charge at pH 7.4, and the GRAVY index^20^, which were computed using BioPython (version 1.77) ProtParam module^21^. We also included the aminoacidic composition. Additionally, codon sequences have been shown to have unexpected effects on the proteins they encode for example, it has been shown they influence local protein structure^22^ and protein lifetime^23^. To evaluate whether the codon sequence influences phosphosite exchange rates, we included the GC content and the content of each codon as features. Finally, we also included protein and peptide turnover rates calculated by Wu *et al.* (2021) on HeLa CCL2 and Kyoto strains^13^.

We extracted features derived from predicted AlphaFold2 protein structures^24^, which were downloaded from AlphaFold DB^25^. Most of these features summarize the structure of the whole protein and the region around the phosphosite (defined as a 10-residue wide window surrounding the phosphosite). They include the secondary structure fractions (both 3-state and 8-state), the median relative surface accessibility (RSA) and the fraction of exposed residues (using an RSA cut-off of ≤0.2) for both the whole protein and the phosphosite region. These features were calculated using DSSP v3.0.0^26^). We also derived features from the predicted local-distance difference test (pLDDT), a per-residue metric of the confidence of the predicted structure, which is also an accurate predictor of protein disorder^27,28^. We included as features the median protein pLDDT, as well as the median pLDDT on a 10-residue wide window surrounding the phosphosite. Finally, we also incorporated features to evaluate the possibility of crosstalk between phosphosites influencing phosphosite exchange rates. This consisted of the number of phosphosites in Ochoa’s *et al.* curated set of phosphosites that are near each phosphosite (taking the phosphosite C*β* as a reference point) and using distance cut-offs of 5, 10 and 15 Å, as well as the distance to the closest annotated phosphosite. Proteins longer than 2700 amino acids are not available whole in AlphaFold DB; they are instead split into multiple predicted structural models due to computational constraints. Because of this, we did not extract features from their predicted structures for such proteins (∼3.3% of proteins in our dataset).

Categorical features were one-hot encoded, while ordinal features were translated into different numerical levels. Constant columns were dropped. This resulted in a total of 945 features.

### Machine learning and phosphosite lifetime prediction

Given that each protein can have multiple phosphosites, and that a small fraction of features contains protein-wide information, simply randomly splitting the dataset into training and test sets could lead to information leakage from the former to the latter. This, in turn, would lead to an overoptimistic estimate of model performance. To avoid this, we cluster proteins by sequence identity, using MMseqs2^29,30^ with greedy incremental clustering (mode 2) and a minimum sequence identity of 30%. The resulting clusters are used to randomly split the dataset into training and test sets (90% and 10% of clusters, respectively), so that proteins belonging to the same cluster are not used in both. Likewise, different cross-validation folds do not contain proteins belonging to the same clusters. Nonetheless, given the large variation in phosphosite turnover within a given protein (Fig. 2b), we expect the effect of any information leakage to be small.

We used gradient boosting trees (as implemented in XGBoost v1.5.2^31^) due to their high generalization potential and ability to incorporate missing data into the training process. The response variable (phosphosite τ) was log10-transformed for training the models. Due to the higher standard error of the measurements for extremely long-lived phosphosites, these were excluded from the dataset (upper bound 10^3.7^ minutes, or ∼83.53 hours). Different model hyperparameters (number of trees, maximum tree depth, learning rate, fraction of features sampled by each tree and fraction of features sampled at each level) were optimized through a grid search by minimizing the squared error. The model was then trained with the estimated optimal parameters and tested on the left-out data.

Feature importance was evaluated by performing SHAP analysis^32^ on the trained model. Based on the game-theoretic concept of Shapley values, SHAP analysis explains each prediction by computing the positive or negative contribution of each feature to the model prediction with respect to the average model output. These values are averaged over the dataset to provide a measure of feature importance. SHAP values were obtained using the TreeExplainer class implemented in SHAP v0.40.0^33^.

### Topic modelling

We used topic modelling, an approach used in natural language processing^34^, to cluster proteins according to their Gene Ontology (GO) terms to ascertain whether phosphorylation exchange rates depend on, for example, the biological processes of the protein or the cellular compartments it belongs to. Additionally, we also used it to study whether these factors influence changes in phosphorylation exchange rates across experimental conditions.

We chose this approach over traditional per-protein GO enrichment analysis for multiple reasons. Each protein can contain multiple phosphosites, and there can be large variation in phosphorylation exchange rates within a given protein (Fig. 2b). Furthermore, distinct phosphosites within a protein could change their phosphorylation exchange rate differently under different conditions. These factors would be difficult to account for with GO enrichment analysis, which we thus find not appropriate for our data. Additionally, topic modelling also has the advantage over regular clustering analysis in that the output indicates which GO terms are distinctive of each group, which is not possible with clustering.

We use non-negative matrix factorization (NMF) as our topic modelling approach to group phosphosites into topics. NMF takes as input a non-negative matrix *V* of dimensions *mxn*, where *m*is the number of phosphosites and *n*the number of GO terms. It then seeks to approximately decompose *V* into two non-negative matrices, the topic-phosphosite matrix *W* and the GO term-topic matrix *H*, such that the dot product of *W* and *H* is approximately equal to *V* . This can be interpreted as finding *p* groups of phosphosites (or “topics”), each of which consists of a weighted combination of the *n* GO terms (defined by the matrix *H*). Conversely, each of the *n* proteins can be understood as a weighted combination of the *p* topics (defined by the matrix *W*). All NMF analyses were performed using using GenSim v4.1.2 available at https://citeseerx.ist.psu.edu/viewdoc/summary?doi=10.1.1.695.4595.

In our approach, each phosphosite is represented as a vector of the different GO terms its protein is annotated with in UniProt^35^. Additionally, we include all of their ancestor terms in the GO ontology. To emphasize more specific terms over more general terms, each element of the *V* matrix is the shortest distance from the corresponding GO term to the top node in the GO ontology. In this way, GO terms are weighted according to their level in the ontology. Ancestral terms and the distance of each term to the top node of the ontology were retrieved using the Python library GOATOOLS v1.2.3^36^.

To ensure convergence of the NMF models, we allowed up to 6000 iterations to learn both the *W* and the *H* matrices. We also set strict convergence thresholds of 10^-6^ to learn both matrices. Topics with probabilities below 0.1 were filtered out. An additional parameter that needs to be set by the user is the number of topics. To determine a good number of topics, we generated NMF models over a range of possible topic numbers (2-50). The choice of the number of topics was guided by the *C_v_* score (which ranges from 0 to 1, from lowest to highest coherence), followed by qualitative human evaluation of the topics generated by NMF models with highest *C_v_* scores (e.g., while a small number of topics may yield a relatively high *C_v_* score, the resulting topics may be hard to interpret, making the model impractical to obtain biological insight). We selected 11 biological process topics (Extended Data Table 1), 8 molecular function topics (Extended Data Table 2) and 12 cellular component topics (Extended Data Table 3). Any other parameters of the model were set to their default values.

Finally, to evaluate whether the phosphosites in each topic have exchange rates that significantly differ from the background (*i.e.*, every other topic), we used the Kolmogorov-Smirnov test, as implemented in SciPy v1.7.3. P-values were adjusted for multiple hypothesis testing using the Benjamini-Hochberg method (FDR < 0.05) using statsmodels v0.13.2 available at https://www.statsmodels.org/.

### KEGG cancer pathways

Through the analyses described above, we identified that phosphosites in signaling-related proteins tend to be shorter lived. In order to showcase the biological and biomedical relevance of this finding, we selected signaling pathways involved in cancer as described in KEGG (KEGG pathway hsa05200). In particular, we selected pathways corresponding to apoptosis, calcium signaling, cell cycle, circadian rhythm, ERK signaling, HIF-1 signaling, JAK-STAT signaling, other RAS signaling, PI3K signaling and WNT signaling. We identified phosphosites involved in these pathways and compared their lifetimes against those of the background (i.e., phosphosites not in proteins within these pathways) using the Kolmogorov-Smirnov test.

### Mean-rank gene-set enrichment

GO terms were assigned to phosphosites using the UniProt identifier of the corresponding protein with the R packages annotationDBI (version 1.62.2) and org.Hs.eg.db (version 3.17.0)^37^. Mean-rank gene-set enrichment was performed to identify terms enriched in highly or lowly ranked turnover phosphosites. Phosphosite turnover values were used with the geneSetTest function from the R package limma (version 3.56.2) (test type = “t”, alternative = “up” or “down”)^38^. Only terms containing at least two quantified phosphoproteins were tested. Within each of the six analysis groups (GO biological process, molecular functions, or cellular compartments enriched in either higher or lower turnover phosphosites), p-values were adjusted by the Benjamini-Hochberg procedure.

### Casein kinase 2 substrate quality

A curated list of 581 human casein kinase 2 (CK2) substrates was extracted from Bradley and collaborators^39^. As in Bradley et al., substrates with S at positions +1 and/or +3 were removed, since these could represent CK2 hierarchical phosphorylation sites which would confound substrate quality analysis. CK2 substrate quality was calculated according to Bradley et al. Briefly, the combined frequencies of D/E at positions -6 to +6 relative to the phosphosite were computed for all filtered CK2 substrates. Then, each substrate was scored by summing the corresponding D/E frequencies for each position that contained D/E and dividing by the maximum possible score (i.e. the summed D/E frequencies across all positions from -6 to +6).

### Associations between kinases and phosphosite exchange rate

We used the data gathered by Bachman et al.^40^ to associate phosphosites to their experimentally determined kinases. We used only associations obtained from PhosphoSitePlus^41^, considered to be the current gold standard. To evaluate whether substrates phosphorylated by a given kinase are associated with lower of higher exchange rates, we performed Kolmogorov-Smirnov tests (using SciPy v1.7.3) of their exchange rates versus the background (i.e., every other phosphosite than is not phosphorylated by that specific kinase). We estimate kinase associations with different exchange by ranking them according to the resulting p-values, similarly to previous work^42^.

### Motif discovery

We set out to identify motifs that could influence phosphosite exchange rates. We extracted 10-residue wide sequence windows surrounding each phosphosite. We subsequently used MEME v5.0.5^43^ to mine motifs in this collection of sub-sequences. We allowed for up to 4 motifs and set a minimum motif width of 5 residues and a maximum width of 10. We used MEME with the zero or one motif occurrence per sequence model option. FIMO 5.0.5^44^ was used to search for individual occurrences of the discovered motifs across the whole dataset using the default settings.

### Details related to the original data in PXD015309

Duplicates of each sample were prepared, one for LC-MS (phosphoproteomics) and one for GC-MS; note that sample 53-58 were exclusively prepared for GC-MS).

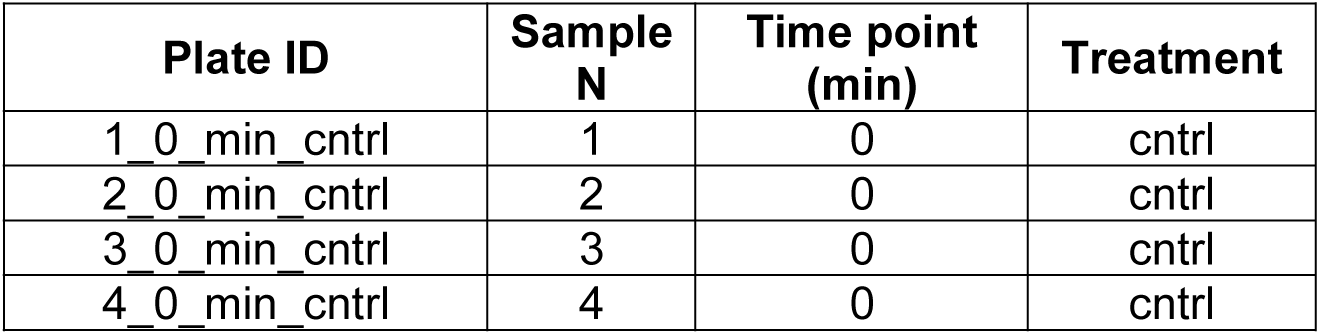
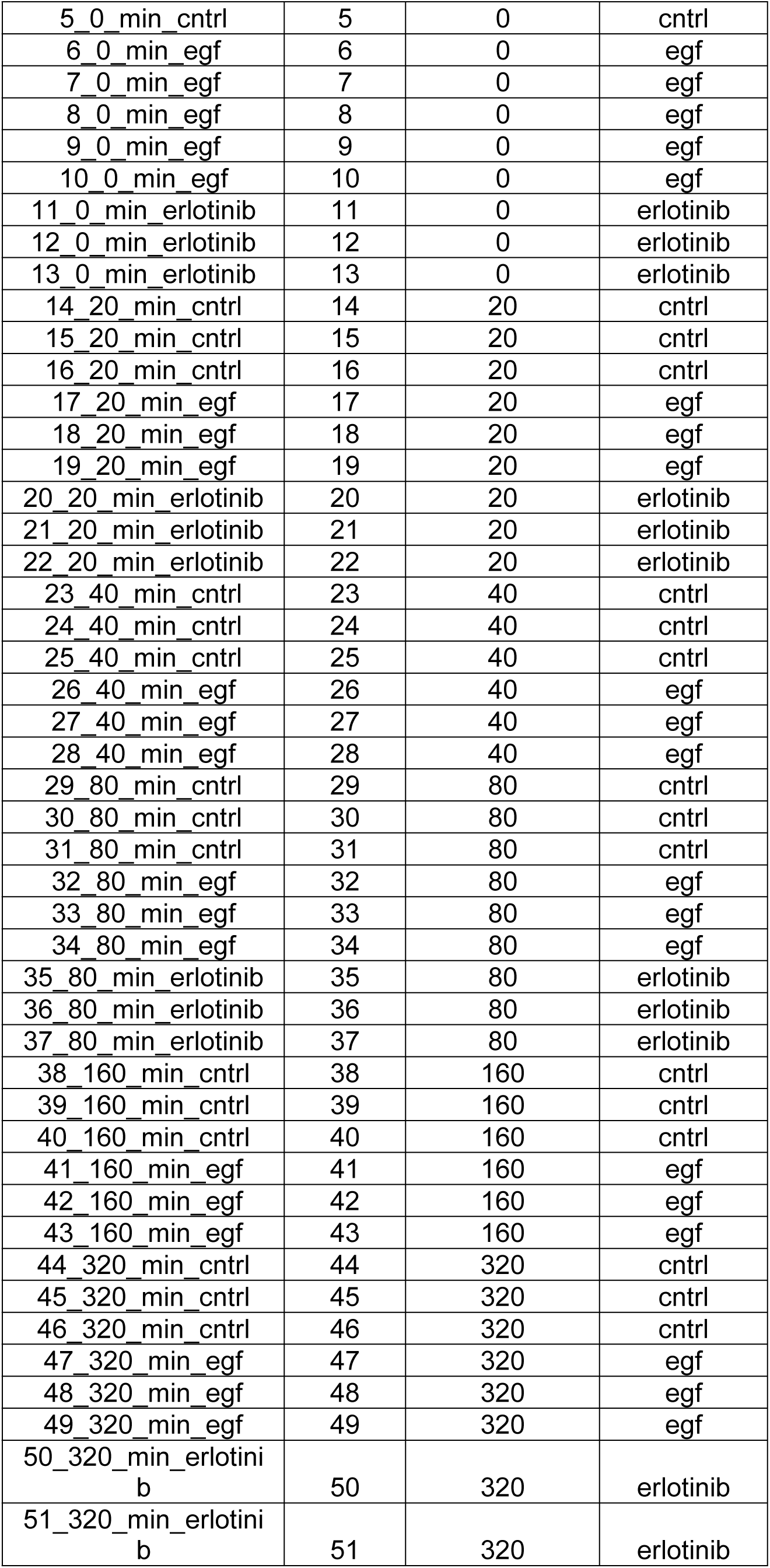
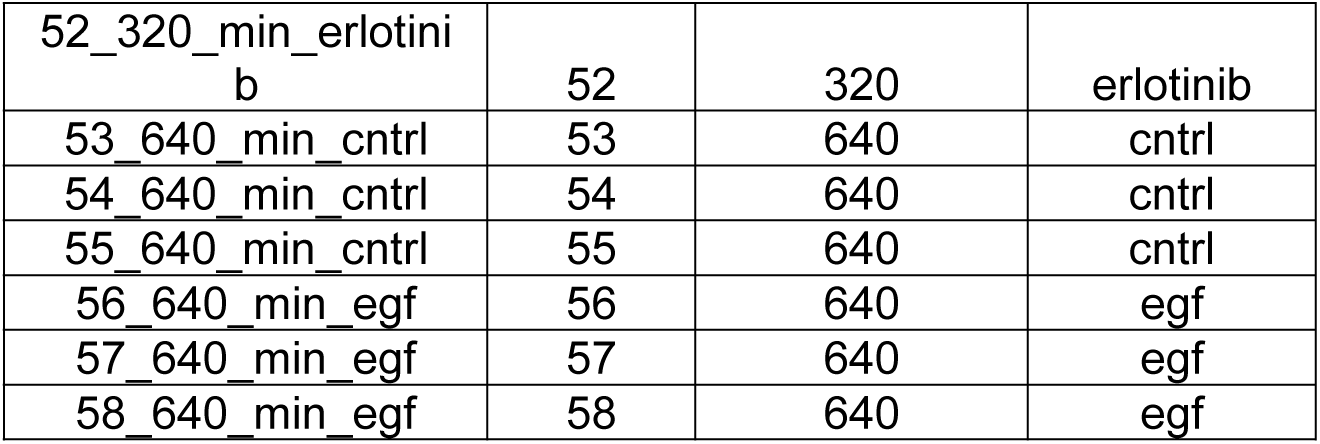

| Plate ID | Sample N | Time point (min) | Treatment |
| --- | --- | --- | --- |
| 1_0_min_cntrl | 1 | 0 | cntrl |
| 2_0_min_cntrl | 2 | 0 | cntrl |
| 3_0_min_cntrl | 3 | 0 | cntrl |
| 4_0_min_cntrl | 4 | 0 | cntrl |
| 5_0_min_cntrl | 5 | 0 | cntrl |
| 6_0_min_egf | 6 | 0 | egf |
| 7_0_min_egf | 7 | 0 | egf |
| 8_0_min_egf | 8 | 0 | egf |
| 9_0_min_egf | 9 | 0 | egf |
| 10_0_min_egf | 10 | 0 | egf |
| 11_0_min_erlotinib | 11 | 0 | erlotinib |
| 12_0_min_erlotinib | 12 | 0 | erlotinib |
| 13_0_min_erlotinib | 13 | 0 | erlotinib |
| 14_20_min_cntrl | 14 | 20 | cntrl |
| 15_20_min_cntrl | 15 | 20 | cntrl |
| 16_20_min_cntrl | 16 | 20 | cntrl |
| 17_20_min_egf | 17 | 20 | egf |
| 18_20_min_egf | 18 | 20 | egf |
| 19_20_min_egf | 19 | 20 | egf |
| 20_20_min_erlotinib | 20 | 20 | erlotinib |
| 21_20_min_erlotinib | 21 | 20 | erlotinib |
| 22_20_min_erlotinib | 22 | 20 | erlotinib |
| 23_40_min_cntrl | 23 | 40 | cntrl |
| 24_40_min_cntrl | 24 | 40 | cntrl |
| 25_40_min_cntrl | 25 | 40 | cntrl |
| 26_40_min_egf | 26 | 40 | egf |
| 27_40_min_egf | 27 | 40 | egf |
| 28_40_min_egf | 28 | 40 | egf |
| 29_80_min_cntrl | 29 | 80 | cntrl |
| 30_80_min_cntrl | 30 | 80 | cntrl |
| 31_80_min_cntrl | 31 | 80 | cntrl |
| 32_80_min_egf | 32 | 80 | egf |
| 33_80_min_egf | 33 | 80 | egf |
| 34_80_min_egf | 34 | 80 | egf |
| 35_80_min_erlotinib | 35 | 80 | erlotinib |
| 36_80_min_erlotinib | 36 | 80 | erlotinib |
| 37_80_min_erlotinib | 37 | 80 | erlotinib |
| 38_160_min_cntrl | 38 | 160 | cntrl |
| 39_160_min_cntrl | 39 | 160 | cntrl |
| 40_160_min_cntrl | 40 | 160 | cntrl |
| 41_160_min_egf | 41 | 160 | egf |
| 42_160_min_egf | 42 | 160 | egf |
| 43_160_min_egf | 43 | 160 | egf |
| 44_320_min_cntrl | 44 | 320 | cntrl |
| 45_320_min_cntrl | 45 | 320 | cntrl |
| 46_320_min_cntrl | 46 | 320 | cntrl |
| 47_320_min_egf | 47 | 320 | egf |
| 48_320_min_egf | 48 | 320 | egf |
| 49_320_min_egf | 49 | 320 | egf |
| 50_320_min_erlotinib | 50 | 320 | erlotinib |
| 51_320_min_erlotinib | 51 | 320 | erlotinib |
| 52_320_min_erlotinib | 52 | 320 | erlotinib |
| 53_640_min_cntrl | 53 | 640 | cntrl |
| 54_640_min_cntrl | 54 | 640 | cntrl |
| 55_640_min_cntrl | 55 | 640 | cntrl |
| 56_640_min_egf | 56 | 640 | egf |
| 57_640_min_egf | 57 | 640 | egf |
| 58_640_min_egf | 58 | 640 | egf |

## Code availability

The scripts and the protocol for determining the phosphopeptide lifetimes (Taus) are currently available for the reviewers on FigShare under:

https://figshare.com/s/e032e659336437d8b059

In this submission a folder with all scripts developed for spectral analysis is included

## Author contributions

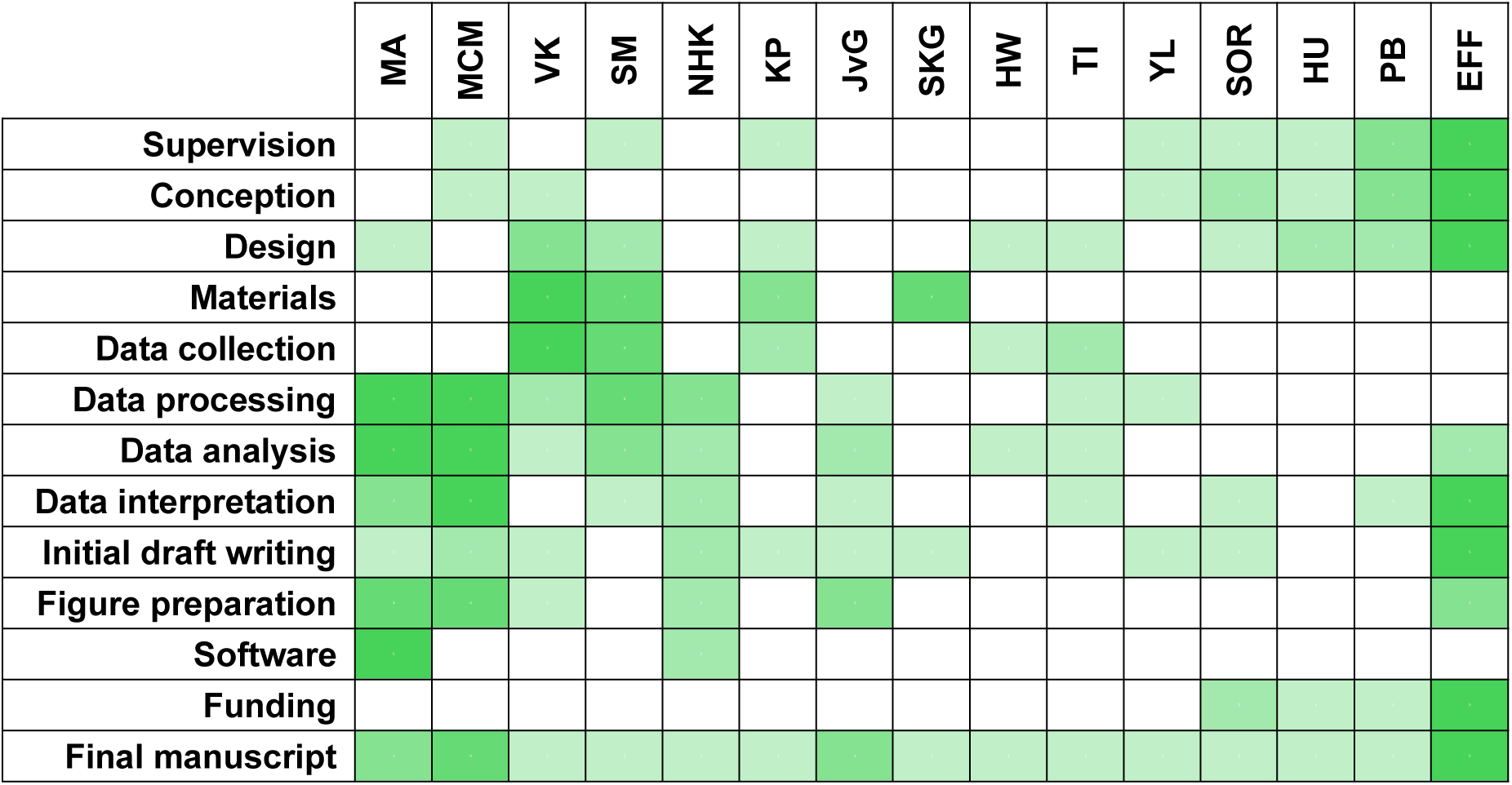

## Funding

EFF was supported by a Schram Stiftung (T0287/35359/2020) and a DFG grant (FO 1342/1-3). EFF also acknowledges the support of the SFB1286, Göttingen, Germany.

## Declaration of competing interests

The authors declare no competing interests.

## Acknowledgments

We thank Janani Durairaj and Mehmet Akdel for their input on the topic modelling analysis, and Nadine Kurz and Youngjun Park for beta testing our code. We would like to also thank Jeffrey N. Savas, Marko Jovanovic, and Ella Doron-Mandel for carefully reading the first version of the manuscript and providing very useful suggestions for improvement. We thank Ivo Feussner, University of Göttingen, for granting access to the GC-MS instrument.

## Notes

### Competing Interest Statement

The authors have declared no competing interest.

