## Extended Data for "Site-specific metabolic labeling reveals proteome-wide phospho-dynamics"

[§] Equally contributing authors

### Extended Data Figures

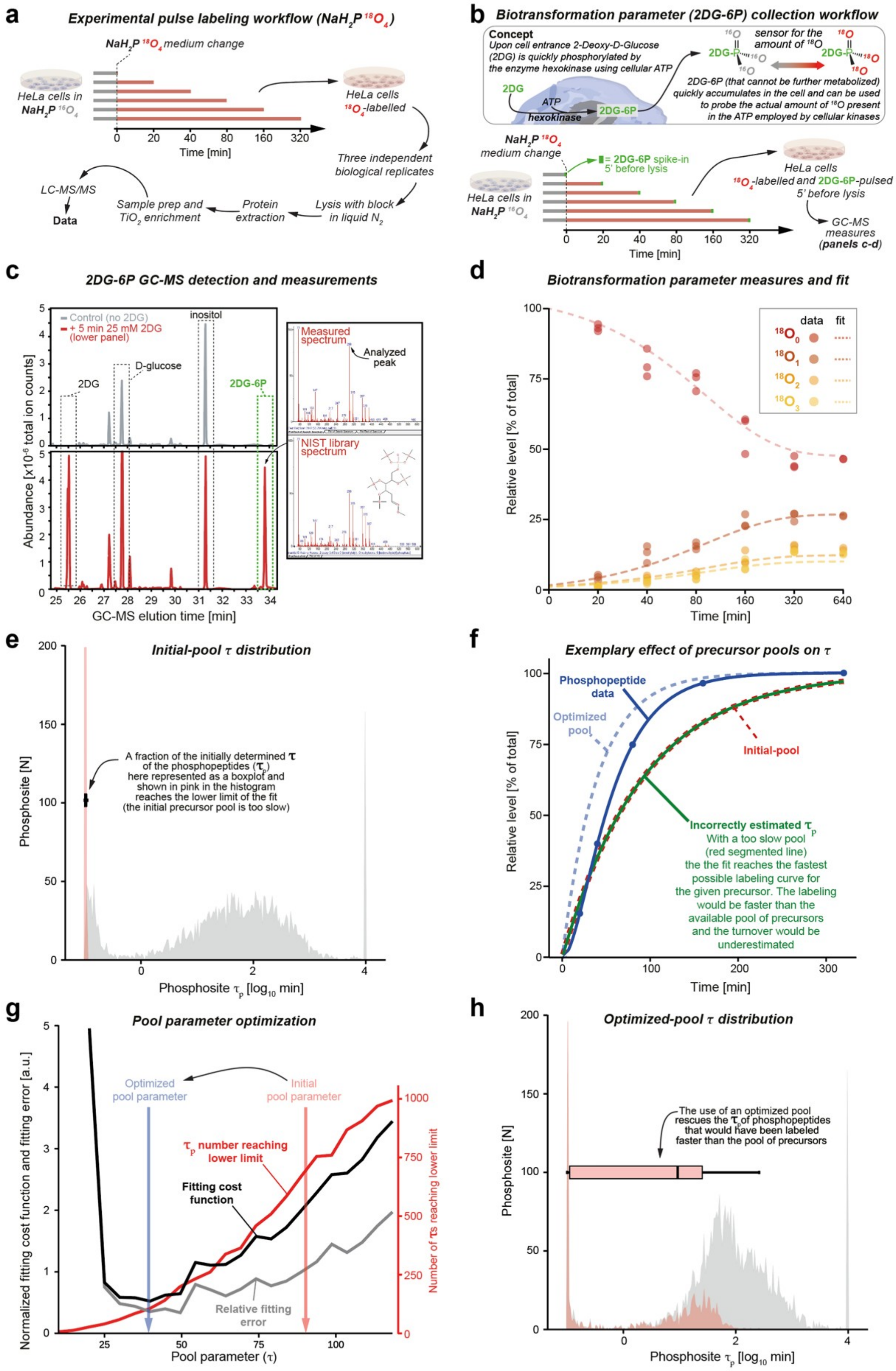


**Extended Data Fig. 1: PulsPhos Experimental workflow and isotopic precursor pool determination.** (**a**) HeLa cells were grown in custom $NaH_2P^{16}O_4$ medium and pulsed for different times in the same medium where $NaH_2P^{16}O_4$ (continues from previous page) was substituted with the stable isotope version ($NaH_2P^{18}O_4$). Three biological replicates were collected for each treatment, quickly lysed in the presence of protease and phosphatase inhibitors, processed with liquid $N_2$ and further processed with standard phosphoproteomics sample preparation (see online methods for details). (**b**) The phosphate group stably labeled with $^{18}O_4$ does not remain as a single unit, but undergoes complex biotransformations before being incorporated into ATP, which is subsequently used by kinases. These biotransformations reduce the amount of $^{18}O$ by diluting it with the $^{16}O$ present in -OH groups that are phosphorylated, thus interfering with the final labeling of the phosphate group in peptides. To determine the amount of $^{18}O$ at each time point and correctly model the precursor pool, we needed to measure it at different times. Since it is not possible to model this "biotransformation parameter", we used an experimental strategy that allowed us to measure it directly inside the cells. The strategy exploits the ability of the hexokinase, an abundant enzyme of the glycolysis that normally catalyzes the phosphorylation of glucose by ATP to glucose-6-P. To "read" the biotransformation parameter at a defined time point and to define the average level of $^{18}O$-labeling of each phosphate group, HeLa cells were normally pulsed in the $NaH_2P^{18}O_4$ medium, but 5 min before lysis, a modified glucose version was added to the culture. This modified glucose (2-deoxy-glucose; 2DG) can be efficiently phosphorylated by the hexokinase generating 2DG-6-Phosphate (2DG-6-P) but cannot be further modified and proceed further in the glycolysis, effectively accumulating in cells after rapid phosphorylation, thereby giving a readout for the labeling status of ATP (**c**) By measuring the amount of $^{18}O$ in 2DG-6P using gas chromatography-mass spectrometry (GC-MS), we were able to accurately determine the amount of $^{18}O$ labeling of phosphate groups at different times (**d**). Note that in GC-MS the 2DG-6P peak is absent when cells are not incubated with 2DG. After 5 min of incubation, the 2DG-6P peak rises rapidly and has very distinct fragmentation peaks that can be used for accurate isotope labeling (inset). (**e**) We extracted the initial pool of precursors using this biotransformation parameter and used it to determine the lifetimes of the phosphopeptides. In doing so, we found that a consistent fraction of the lifetimes overlapped with the shorter limit and appeared close to zero (red box-plot in the left corner of the image). This is likely due to the fact that the biotransformation parameter measured with the hexokinase strategy averages the pool over the 5 minutes required to obtain a sufficient 2DG-6-P peak, thus the speed of the measured pool is perceived as slower. At the same time, when using shorter 2DG pulses, not enough signal was available to determine the isotopic composition of the oxygen within its phosphate group in GC-MS, so we maintained the 5 min spike-in time window (data not shown). (**f**) To understand this from another perspective, a too-slow estimation of the precursor pool (red segmented line) would make fast phosphopeptides appear being labelled faster than the pool itself (green continuous line), leading to a too-fast estimation of $\tau_{\mathrm{p}}$ (green line), even reaching the lower fitting limit, as seen in the “too-fast” fraction in e. To "rescue" the too short $\tau_{\mathrm{p}}$, we followed a pool optimization strategy, assuming that a precursor pool with a faster kinetic might exist (light blue segmented line) and would obviously be labeled faster than the available $^{18}O$-label that could be incorporated into phosphopeptides (darker blue line). (**g**) This strategy was based on minimizing a combined cost function consisting of the relative fitting error and the relative number of “too-short” lifetimes (i.e. reaching the lower fitting limit), leading to the determination of an optimized kinetic pool parameter ($\tau$); see online methods for details. (**h**) Indeed, this pool optimization strategy allowed to determine several lifetimes that would otherwise have been labeled faster than the pool of precursors (note the red-box plot in the upper part of the figure that moves on the right side of the distribution compared to panel e).

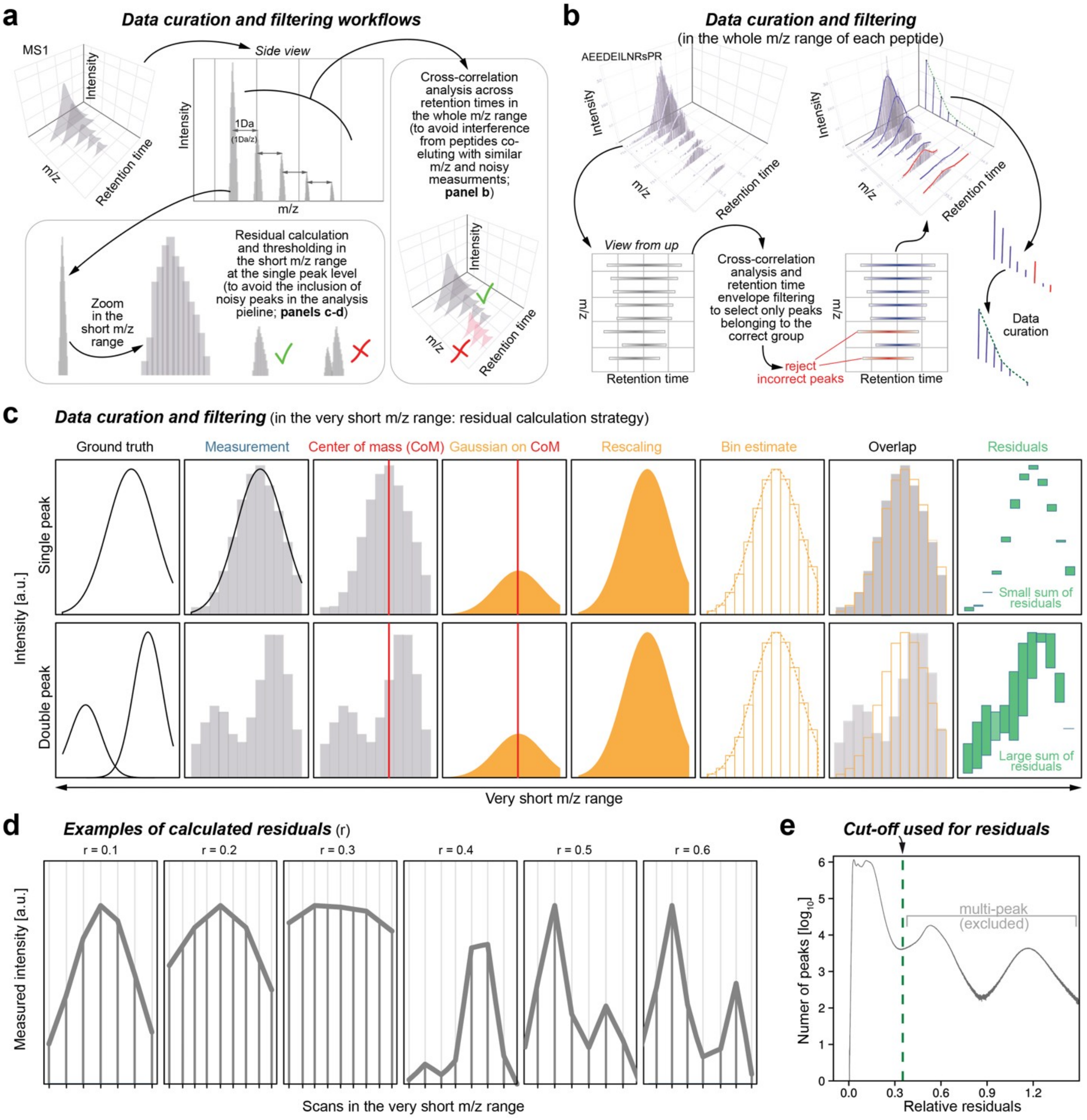


**Extended Data Fig. 2: PulsPhos data curation and filtering strategies.** (**a**) Schematic representation of the two aspects considered in our data curation and filtering strategy: consistency of the poly-isotopic peaks across the MS1 m/z range for each phosphopeptide (explained in panel b) and peak sharpness and reliability in the very short m/z range (corresponding to a single isotopic peak; explained in panels c-d). We have found that our method is very sensitive to contamination and poor quality measurements in MS1, and the data curation and filtering step is important to ensure correct lifetime estimation. We are aware that more sensitive and higher resolution instrumentation will further improve the coverage that our method can achieve in the future. (**b**) Throughout the m/z range of each peptide, contaminants that do not pass the cross-correlation envelope analysis (which can be identified by differences in retention time as shown in the example) are not considered for data integration. (**c**) Similarly, even in the very short m/z range, we have found that poorly measured peaks do not have a clear single peak distribution, but rather a multi-peak distribution. We evaluate this by measuring the residuals following a Gaussian fitting strategy. Multi-peak distributions (lower row of the two exemplary sets) have a large sum of residuals, as also shown in panel **d**. We set up a cut-off residual sum from perfect gaussians, shown in **e**, that excludes the majority of multi-peak distributions and preserves the reliability of MS1 measurements in the short m/z range, ensuring the quality of the data used for our lifetime estimations.

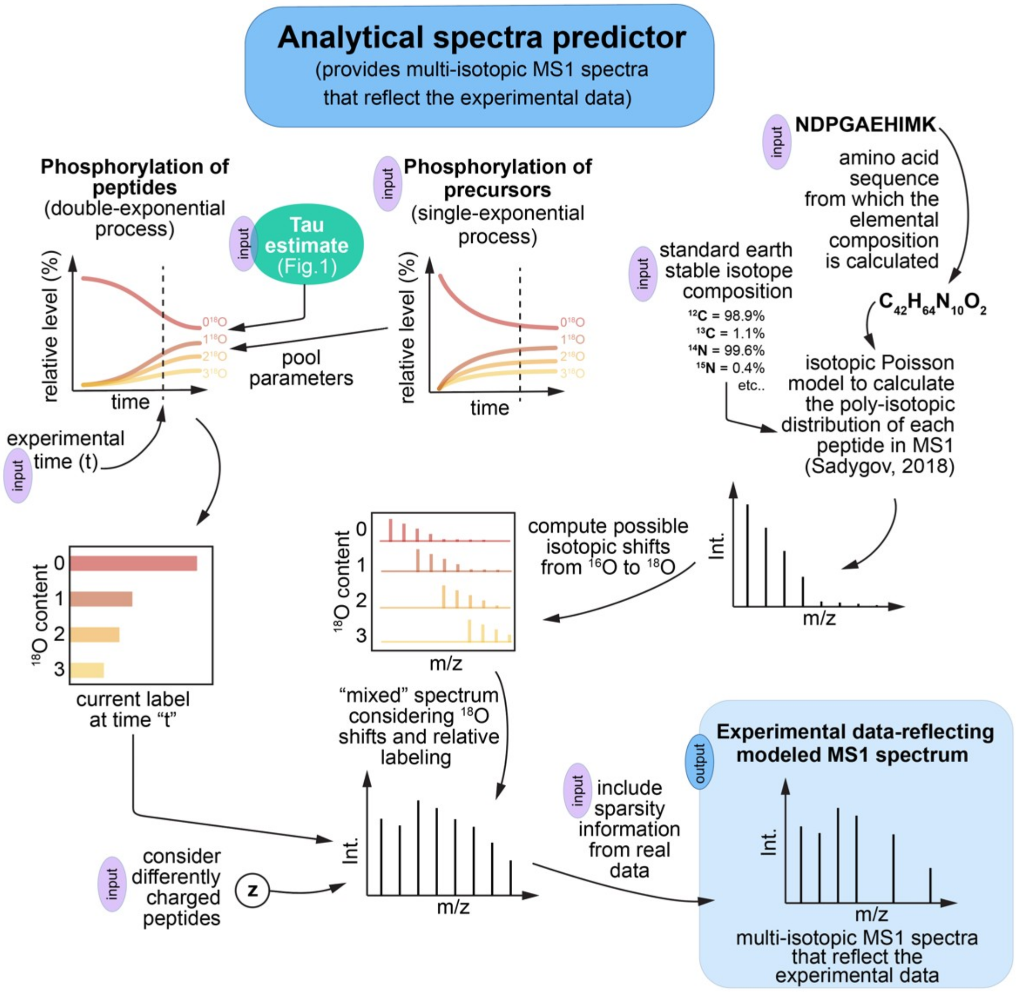


**Extended Data Fig. 3: Analytical spectral predictor pipeline.** The approach used for the fitting requires to model MS1 spectra that reflect multi-isotopic spectra close to the experimental data (also referred to as "in silico peptide spectral families" as defined in main Fig. 1b). Our spectral predictor pipeline requires seven inputs to be able to produce multi-isotopic MS1 spectra that closely reflect the experimental data. These are used at later steps for model-data comparisons that allow the precise determination of the phospho-lifetime (main Fig. 1b). The 7 inputs include: (1) The amino acidic sequence of the phosphopeptide and (2) The standard earth isotope composition, that are necessary to simulate the poly-isotopic distribution of each peptide in MS1; (3) The phosphorylation of the precursor phosphate groups as determined in Ext. Data Fig. 1d. The precursor pools are fit to the measured data to get the curve parameters. With these pool parameters as input and in combination with the tau and the labeling time, the phospho labeling for this time is calculated with a two-exponential curve. (4) The Tau estimate (Main Fig. 1b) and (5) The labelling time, which allow to determine the labeling extent at the defined time "t". In the modeling, also the differently charged peptides (6) and the information sparsity specific for measured data (7) are considered. Altogether these seven inputs allow to predict reliable *in silico* peptide families that can be used for lifetime determination. The total difference between all measured and theoretical spectra for all measurements, weighted by detector counts of a measurement, is used as cost function to optimize the single fitting turnover parameter.

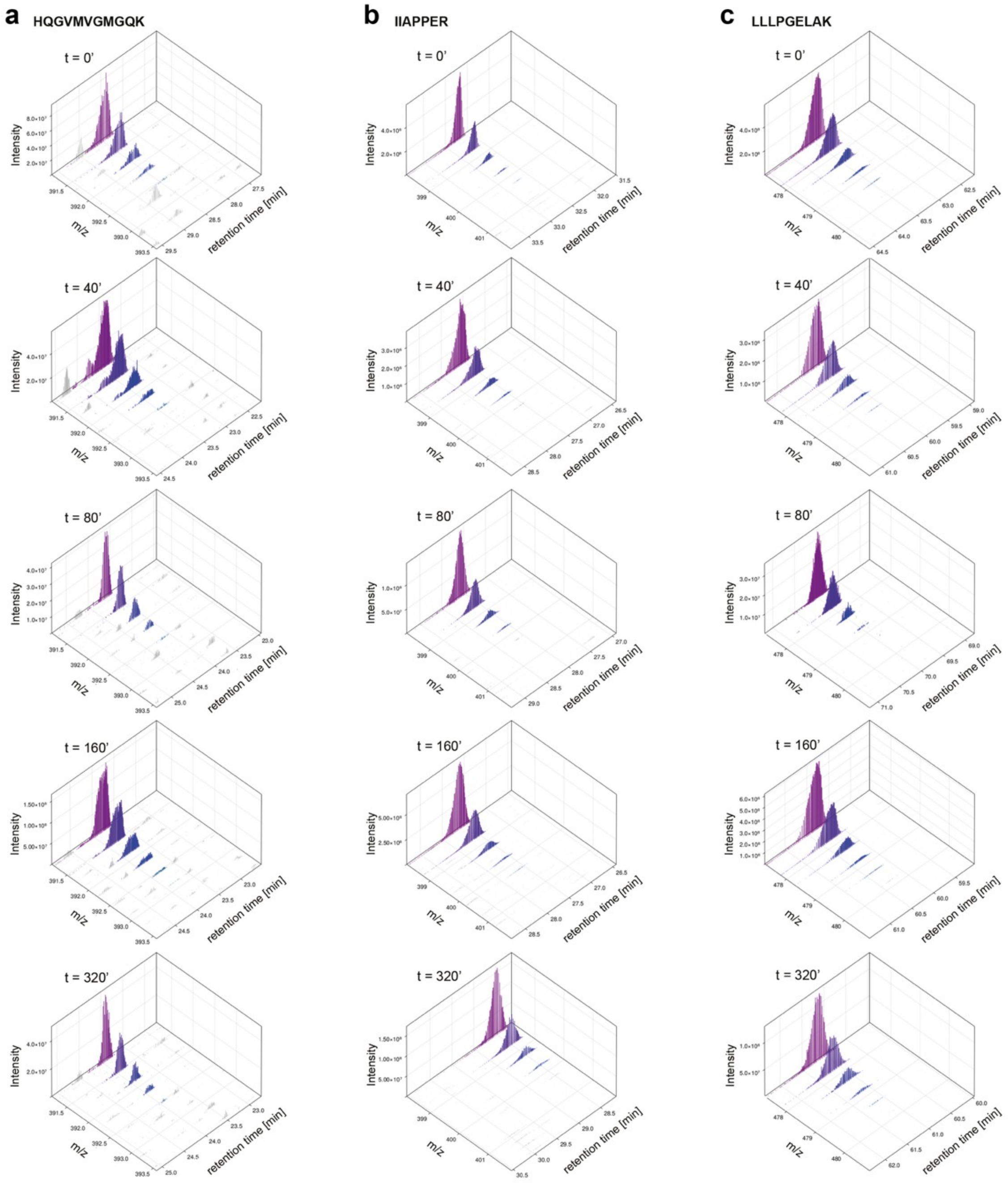


**Extended Data Fig. 4: Non-phosphorylated peptide examples show no $^{18}O$ labeling within the considered labeling time frame.** (**a-c**) Three examples of MS1 labeling time courses showing that the structural labeling due to $^{18}O$-labeling introduced by the $^{18}O$-labeled phosphate groups can be neglected within the short labeling pulses used in this work. In the phosphopeptide enriched samples we do not see many unphosphorylated peptides across different pulse times (~300). In any case, none of those that we detected shows a labeling shift as we can clearly observe in phosphopeptides (see as an example the shift towards higher m/z for the phosphopeptide shown in main Fig. 1, panel c). We run this control to exclude that the $^{18}O$ labeling could lead to an m/z shift of our peptides not because of a *bona-fide* phosphate group labeling but due to an unspecific $^{18}O$ labeling of other oxygens present in these peptides. It also should be noted that the amount of $^{18}O$ introduced with phosphate groups is extremely small when compared to the endogenous $^{16}O$ present in cells, which probably serves as a large buffer for the $^{18}O$ and makes possible unspecific labeling of the backbone and the OH groups within peptides neglectable.

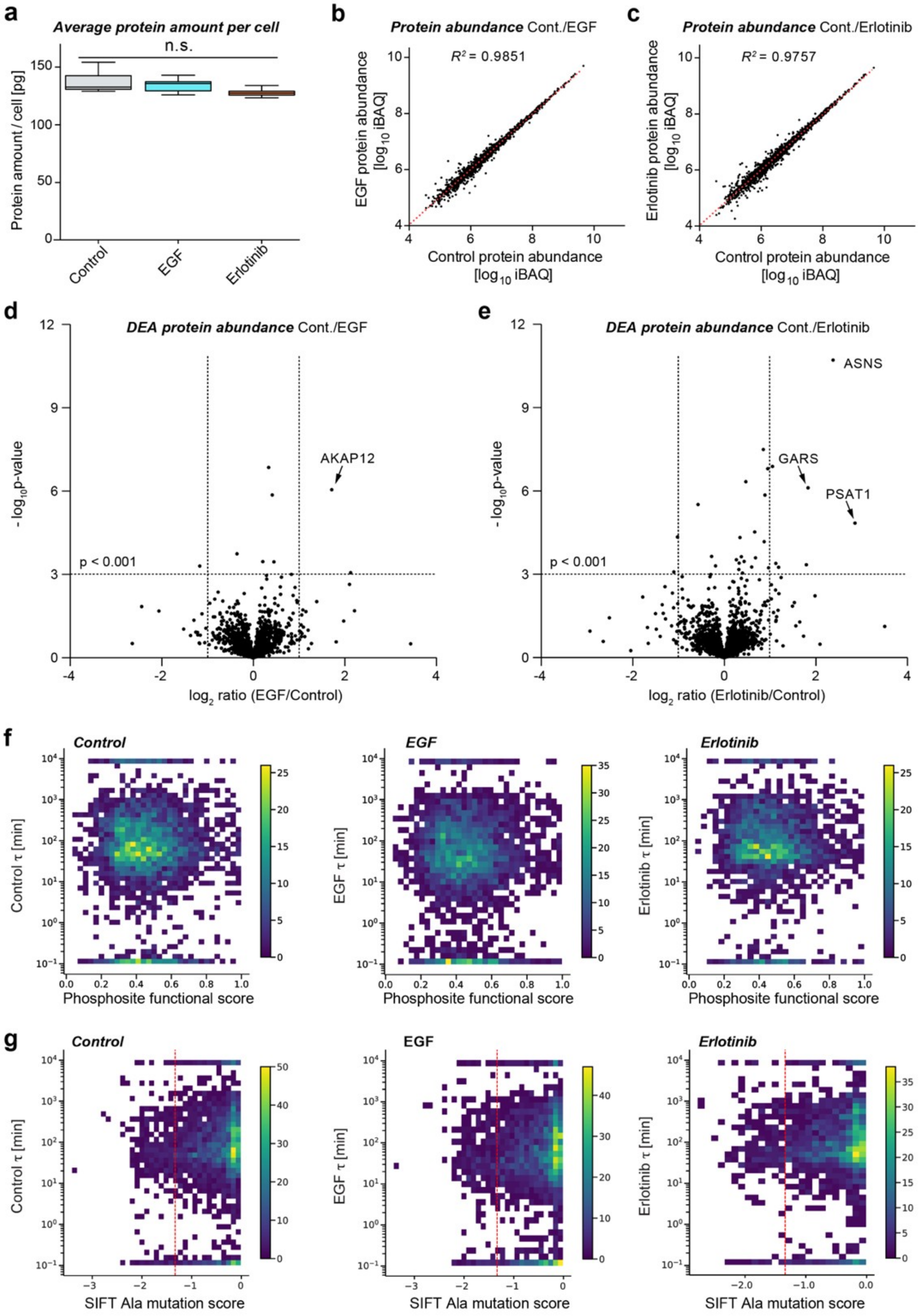


**Extended Data Fig. 5: Pharmacological treatments do not affect main proteome composition and phosphopeptide turnover properties.** (**a**) Estimated protein amount per cell. (**b-c**) Scatter plot of protein abundance showing the comparison between control and EGF (b) or control and Erlotinib treatment (c). (**d-e**) Volcano plots showing differential protein abundance analysis for the control and EGF (d) or control and Erlotinib treatment (e). Note minimal changes. (**f**) Functional score matrix vs the measured lifetimes, showing little correlation for all for the three samples analyzed, similarly to (**g**) showing the log10-transformed SIFT Ala mutation score. The latter predicts whether an amino acid substitution affects protein function and ranges from 0.0 (deleterious) to 1.0 (tolerated).. The red line indicates the threshold (0.05) below which variants are considered deleterious.

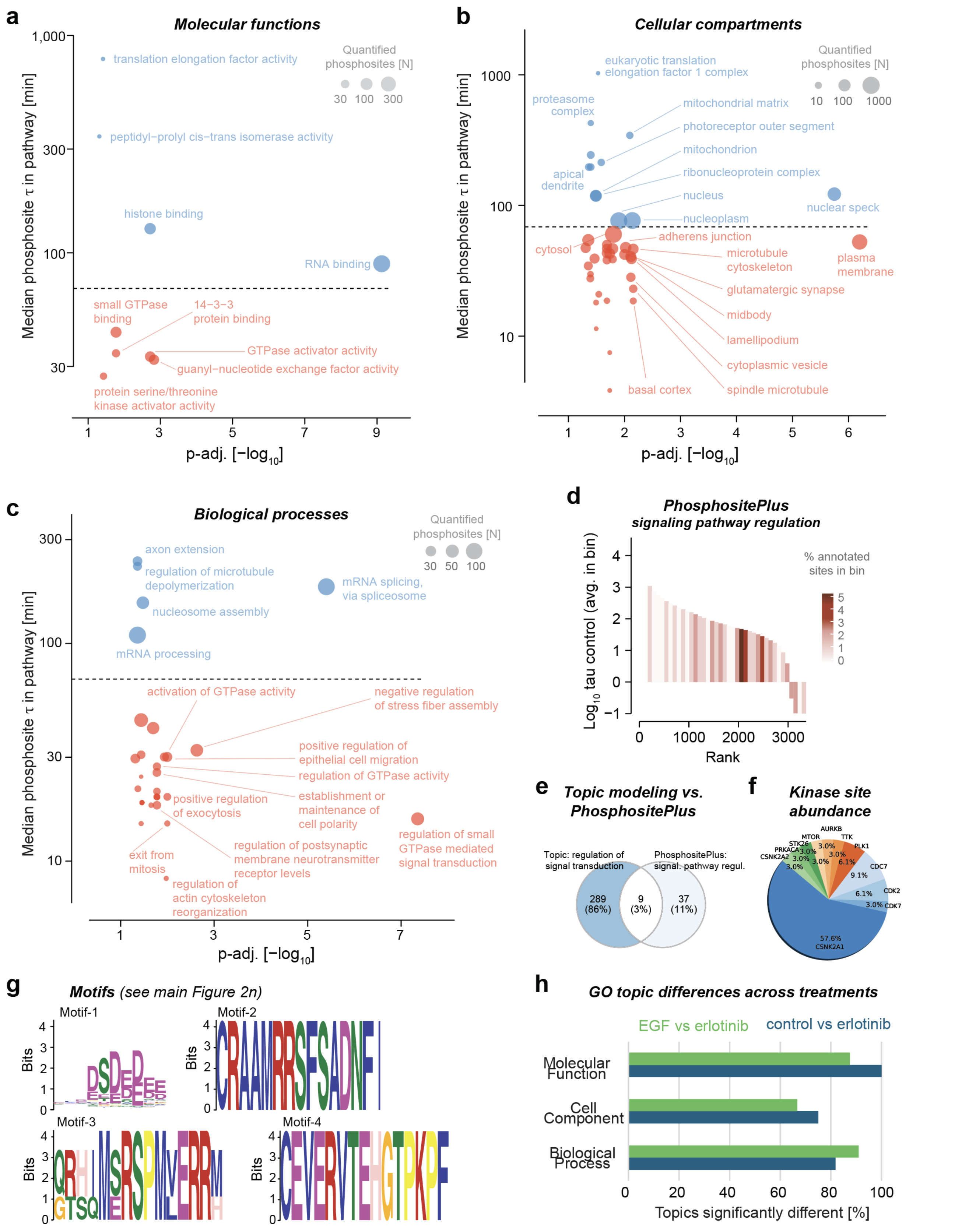


**Extended Data Fig. 6: Gene ontology and phosphorylated motif analysis.** (**a-c**) Mean-rank gene set test analysis for the untreated samples largely recapitulates what observed in the topic modeling analysis (main Fig. 1 h-j). Only GO (continues from previous page) terms significantly enriched in either highly or lowly ranked phosphosites (adjusted p-value < 0.05) are shown and the top 10 most significantly enriched terms are labelled. Blue color indicates significantly slower exchange rates and red significantly faster(**d**) Phosphosites annotated in PhosphositePlus to the biological process "signaling pathway regulation" show a negative correlation with the measured lifetimes, indicating that phosphorylation events linked to signaling pathway regulation, which probably requires fast responses in cells are indeed associated to short living phosphorylation events. Note that this result is similar to what we have observed in our topic modeling analysis (main Fig. 2j) although, as we show in panel **e**, it is not redundant as the phosphosites in panel **d** and those in the modeled topic in Fig. 2j overlap little. (**f**) Abundance of the sites recognized by different kinases in our dataset. (**g**) Significant motifs found by motif analysis (with >1 peptide), referred to main Fig. 2n. (**h**) Significantly different topics ($p<0.05$) across treatments upon percentile normalization. Significant differences between WT and EGF were not found upon percentile normalization (not shown). Overall, after percentile normalization, lifetimes increase upon erlotinib treatment, suggesting that blocking EGF signaling in HeLa cells has a profound effect on phosphorylation dynamics.

| Topic summary | Top 3 GO terms | Coherence score ($C_v$) |
| --- | --- | --- |
| Regulation of cytoskeletal organization | Regulation of cytoskeleton organization<br>Regulation of protein polymerization<br>Regulation of supramolecular fiber organization | 0.54 |
| Regulation of signal transduction | Positive regulation of cellular processes<br>Regulation of intracellular signal transduction<br>Positive regulation of biological process | 0.41 |
| Organelle organization & protein localization | Cellular component assembly<br>Cellular component organization<br>Organelle organization | 0.37 |
| Protein ubiquitination | Protein ubiquitination<br>Protein modification by small protein conjugation<br>Protein modification by small protein conjugation or removal | 0.85 |
| Transcriptional regulation | Regulation of nucleic acid-templated transcription<br>Regulation of RNA biosynthetic process<br>Regulation of transcription, DNA-templated | 0.82 |
| Negative transcriptional regulation | Negative regulation of nucleic acid-templated transcription<br>Negative regulation of macromolecule biosynthetic process<br>Negative regulation of cellular biosynthetic process | 0.89 |
| RNA processing | Nucleic acid metabolic process<br>Macromolecule metabolic process<br>RNA processing | 0.82 |
| Positive transcriptional regulation | Positive regulation of nitrogen compound metabolic process<br>Positive regulation of macromolecule metabolic process<br>Positive regulation of nucleic acid-templated transcription | 0.68 |
| Negative regulation of phosphorylation | Negative regulation of phosphorylation<br>Negative regulation of protein phosphorylation<br>Negative regulation of phosphate metabolic process | 0.42 |
| Negative regulation of apoptosis & cell communication | Negative regulation of apoptotic process<br>Negative regulation of cell communication<br>Negative regulation of programmed cell death | 0.58 |
| Regulation of cell projection organization | Regulation of plasma membrane bounded cell projection organization<br>Regulation of neuron projection development<br>Regulation of cellular component organization | 0.46 |

**Extended Data Table 1:** Biological process topics,

| Topic summary | Top 3 GO terms | Coherence score ($C_v$) |
|---|---|---|
| ATP binding & kinase activity | ATP binding<br>Adenyl ribonucleotide binding<br>Adenyl nucleotide binding | 0.71 |
| Nucleic acid binding | Nucleic acid binding<br>RNA binding<br>DNA binding | 0.49 |
| RNA binding & pyrophosphatase activity | Nucleoside-triphosphatase activity<br>RNA binding<br>Pyrophosphatase activity | 0.62 |
| ATP hydrolysis | ATP hydrolysis activity<br>ATP binding<br>Adenyl nucleotide binding | 0.67 |
| Transcription factor & RNA polymerase II binding | Enzyme binding<br>DNA-binding transcription factor binding<br>Protein binding | 0.65 |
| Cytoskeletal protein binding | Cytoskeletal protein binding<br>Metal ion binding<br>RNA polymerase II-specific DNA-binding transcription factor binding | 0.35 |
| Protein kinase binding & kinase activity | Protein kinase binding<br>Adenyl nucleotide binding<br>ATP binding | 0.68 |
| Cadherin binding | Cadherin binding<br>Metal ion binding<br>Ion binding | 0.60 |

**Extended Data Table 2:** Molecular function topics.

| Topic summary | Top 3 GO terms | Coherence score ($C_v$) |
|---|---|---|
| Polymeric cytoskeletal fiber | Polymeric cytoskeletal fiber<br>Supramolecular fiber<br>Microtubule | 0.36 |
| Cytoplasmic vesicle | Cytoplasmic vesicle<br>Intracellular vesicle<br>Intracellular non-membrane-bounded organelle | 0.33 |
| Nucleus | Nucleus<br>Intracellular membrane-bounded organelle<br>Intracellular organelle | 0.41 |
| Nucleus & extracellular vesicles | Nucleus<br>Intracellular membrane-bounded organelle<br>Membrane-bounded organelle | 0.45 |
| Intracellular organelles & exosomes | Intracellular membrane-bounded organelle<br>Extracellular exosome<br>Membrane-bounded organelle | 0.42 |
| Microtubules & cell-cell junctions | Microtubule<br>Polymeric cytoskeletal fiber<br>Supramolecular fiber | 0.47 |
| Nucleus & membrane | Nucleus<br>Intracellular membrane-bounded organelle<br>Membrane | 0.37 |
| Cytoplasmic vesicles | Cytoplasmic vesicle membrane<br>Bounding membrane of organelle<br>Vesicle membrane | 0.45 |
| Cell-substrate junctions | Focal adhesion<br>Cell-substrate junction<br>Secretory granule lumen | 0.63 |
| Endosome | Endosome<br>Cytoplasmic vesicle membrane<br>Synaptic vesicle | 0.64 |
| Cytoskeleton | Cytoskeleton<br>Intracellular non-membrane-bounded organelle<br>Non-membrane-bounded organelle | 0.50 |
| Organelle membrane | Organelle membrane<br>Bounding membrane of organelle<br>Lysosomal membrane | 0.27 |

**Extended Data Table 3:** Cellular component topics.

### A practical guide for the scripts used for the determination of Phospho-lifetimes

In order to run the scripts, you will require a computer that has at least 32GB RAM and 1TB of hard disk space. In our experience this works best in a high-performance computing server.

This practical guide has been optimized for Ubuntu 22.04.3 LTS https://ubuntu.com/download/desktop/thank-you?version=22.04.3&architecture=amd64
For minimal support in running and installing Ubuntu please refer to https://ubuntu.com/tutorials/install-ubuntu-desktop#1-overview

Before starting it will be necessary to download the directory called "data", containing the .mz5 and the experimental information using the following path and the password:

**Path:** https://owncloud.gwdg.de/index.php/s/LWmAxgwqw0YGxX0/authenticate
**Password:** Phospho

Ideally this directory should be placed in the "Downloads" folder if in the local computer (~/Downloads/)

Note that you will need ~150GB to run the whole directory.

Your data directory should be structured as follows

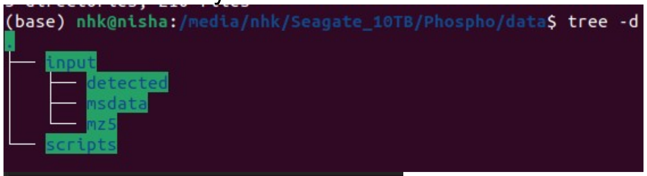


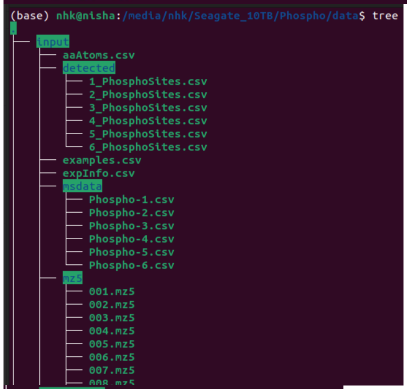


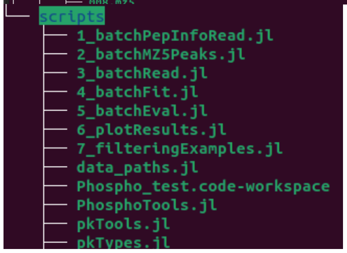

### Detailed instructions for the general setup and analysis in Julia:

**1. Install Julia in Ubuntu through the command line:**
Open the terminal and then copy the following command:
`sudo snap install julia –classic`

If you do not have sudo (root / administrator) rights copy the following command to the terminal
`wget -v " https://julialang-s3.julialang.org/bin/linux/x64/1.9/julia-1.9.2-linux-x86_64.tar.gz"`

Note that in our experience Julia version 1.9.2 works best and has all the packages
For troubleshooting Julia installation please refer to: https://julialang.org/downloads/

**2. In Julia, install all the necessary packages:**

- Type `julia` in the command line to start Julia
- Type `]` in Julia to prompt the installation of the packages
- Type the following line that will install all the necessary packages:
  `add CSV, HDF5, SpecialFunctions, StatsBase, ProgressMeter, Statistics, Distributions, Optim, ForwardDiff, DataFrames, Glob, JLD2, Printf, CairoMakie`

The installation will require some seconds and at the end of it you should see something like

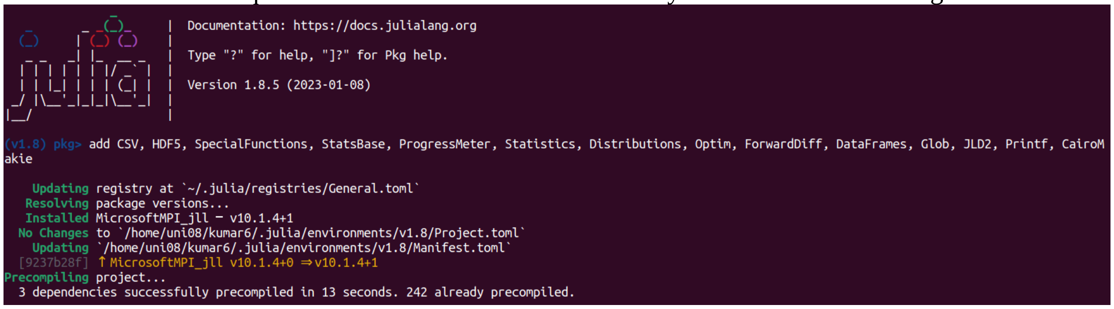


**3. Exit from Julia**
Please exit from Julia by pressing ctrl+z

**4. Check the data paths:**
If the script is run in the local computer and is found in the following path (~Downloads/data/) you can directly move to the next step.

If the data folder is in a different path you will need to edit the script.

You will find the script file in the downloaded folder (/data/scripts/data_paths.jl). To edit it in line 4 adjust the path in accordance to where you located the downloaded folder `(cfg["path_phospho"]=...)`

**5. Move to the "scripts" directory in the Ubuntu terminal**
If downloaded in the downloads folder, then move to the downloads folder using the following command in the terminal `cd /Download/data/scripts/`

If it is not in that folder use the appropriate path in the `cd` command

**6. Run the first script to import all the tools**

- First enter Julia by typing `julia` in the command line
- To run the script in Julia type `include(PhosphoTools.jl)` and hit enter

Note that this script should run quickly (usually less than a minute)
**7. Run the remaining scripts (1 to 7)**

By default, the scripts are commented in a way that will allow to quickly analyze only the first .mz5 for testing purposes. If you want to run the whole 180 files please refer to the note at the end of this point.
For each of the scripts follow the subsequent set of instructions in Julia:

- Type `include(/path/on/server/scripts/1_batchPepInfoRead.jl)` and hit enter
- Wait until the script is finished (you will see the Julia logo in the command line)
- Type `include(/path/on/server/scripts/2_batchMZ5Peaks.jl)` and hit enter
- Wait until the script is finished (note that this step might take some hours)
- Type `include(/path/on/server/scripts/3_batchRead.jl)` and hit enter
- Wait until the script is finished (this step should be faster than the previous one)
- Type `include(/path/on/server/scripts/4_batchFit.jl)` and hit enter
- Wait until the script is finished (you will see the Julia logo in the command line)
- Type `include(/path/on/server/scripts/5_batchEval.jl)` and hit enter
- Wait until the script is finished (you will see the Julia logo in the command line)
- Type `include(/path/on/server/scripts/6_plotResults.jl)` and hit enter
- Wait until the script is finished (you will see the Julia logo in the command line)
- Type `include(/path/on/server/scripts/7_filteringExamples.jl)` and hit enter
- Wait until the script is finished (you will see the Julia logo in the command line)

*****Note that it may happen that Julia crashes during a complex computation.** In that case, one can skip to a specific script but Phosphotools.jl <u>must always be run first.</u>

Note that for running all the 180 files you will need to edit all the scripts manually by going to the end of each one of them and uncommenting the line for the 180 files (see following images)

Initial settings (for running just the first .mz5 file) eg: `2_batchMZ5Peaks.jl`

```
# 1. batch process of mz5 files
#
batchPeaks(cfg, [1])                                 # test: for 1st file
# batchPeaks(cfg, 1:180)                               #       for all 180 files (can take hours)
```

Modified settings for running the 180 files

```
# 1. batch process of mz5 files
#
#batchPeaks(cfg, [1])                                 # test: for 1st file
 batchPeaks(cfg, 1:180)                               #       for all 180 files (can take hours)
```

**8. Check the output**
If the scripts run without problems, you should find the output files in the “results” folder

```
.
├── chunks
├── fig
├── fit
├── peaks
└── pic
    ├── AQQELVPPQQQAsPPQLPK
    │   └── cntrl
    ├── DLVQPDKPAsPK
    │   └── cntrl
    ├── GLGHPPSsPLLK
    │   └── cntrl
    ├── ILKsPEIQR
    │   ├── cntrl
    │   └── egf
    ├── MLPHAPGVQMQAIPEDAVHEDsGDEDGEDPDKR
    │   └── cntrl
    ├── RKsEPAVGPPR
    │   └── cntrl
    └── RRDsDGVDGFEAEGK
        └── cntrl
```
